# L(3)mbt and the LINT complex safeguard tissue identity in the Drosophila ovary

**DOI:** 10.1101/178194

**Authors:** Rémi-Xavier Coux, Felipe Karam Teixeira, Ruth Lehmann

**Affiliations:** Howard Hughes Medical Institute (HHMI) and Kimmel Center for Biology and Medicine of the Skirball Institute, Department of Cell Biology, New York University School of Medicine, New York, NY 10016, USA; Department of Genetics, University of Cambridge, Downing Street, Cambridge CB2 3EH, UK

**Keywords:** L(3)mbt, tissue identity, Nanos, oogenesis, Drosophila, Lint1

## Abstract

Maintenance of cellular identity is essential for tissue development and homeostasis. At the molecular level, cell identity is determined by the coordinated activation and repression of defined sets of genes. Defects in the maintenance of the genetic programs required for identity can have dire consequences such as organ malformation and cancer. The tumor suppressor L(3)mbt was shown to secure cellular identity in Drosophila larval brains by repressing germline-specific genes. Here we interrogate the temporal and spatial requirements for L(3)mbt in the Drosophila ovary, and show that it safeguards the integrity of both somatic and germline tissues. *L(3)mbt* mutant ovaries exhibit multiple developmental defects, which we find to be largely caused by the inappropriate expression of a single gene, *nanos*, a key regulator of germline fate, in the somatic cells of the ovary. In the female germline, we find that L(3)mbt represses testis-specific and neuronal genes. Molecularly, we show that L(3)mbt function in the ovary is mediated through its cofactor Lint1 but independent of the dREAM complex. Together, our work uncovers a more complex role for L(3)mbt than previously understood and demonstrates that L(3)mbt secures tissue identity by preventing the simultaneous expression of original identity markers and tissue-specific misexpression signatures.

## Introduction

Development requires tight control of gene expression as differentiating cells must express lineage-specific genes while repressing genes that promote other fates. Mechanisms ensuring the maintenance of cellular identity must be robust, as changes in cell fate appear to be fairly uncommon in wild-type conditions, with only one documented case of regulated, complete fate switch being described in *C. elegans* (Jarriault et al., 2008). However, rare cases of transdifferentiation have been observed in mutants affecting chromatin complexes, suggesting a role for chromatin structure in the maintenance of cellular identity (Petrella et al., 2011; Tursun et al., 2011). A notable example is given by mutations affecting the Drosophila *lethal (3) malignant brain tumor* (*l(3)mbt*) gene, which cause malignant brain tumors that ectopically express germline-specific genes and have been proposed to be soma-to-germline transformations (Gateff et al., 1993; Janic et al., 2010). Gene expression profiling of *l(3)mbt* brain tumors and L(3)mbt-depleted cultured somatic cells identified a group of upregulated genes known as the Malignant Brain Tumor Signature (MBTS) that is enriched for factors specifically expressed in germ cells (Georlette et al., 2007; Janic et al., 2010; Meier et al., 2012; Sumiyoshi et al., 2016). Mutations of germline-specific genes, including those impairing the piRNA factors *piwi*, *aub*, and *vasa*, as well as the translational repressor *nanos*, were found to suppress the neural overgrowth induced by loss of L(3)mbt (Janic et al., 2010). A subsequent study provided evidence that the Hippo growth control pathway is critical for *l(3)mbt* mutant brain overgrowth, suggesting an alternative cause of tumorigenesis (Richter et al., 2011). Furthermore, our lab showed that strong *l(3)mbt* mutations cause a maternal, germline autonomous phenotype that precludes normal embryonic development including primordial germ cell formation (Yohn et al., 2003). Together, these studies suggest that L(3)mbt may impart many functions in regulation of tissue identity.

*L(3)mbt* encodes a 1477 amino-acid protein that is ubiquitously expressed in Drosophila and is conserved from worms to humans. L(3)mbt is thought to be a chromatin reader and harbors three MBT repeats that bind methylated histone tails *in vitro* as well as a Zinc-Finger domain (Bonasio et al., 2010). L(3)mbt is enriched at the promoters of repressed genes, suggesting a direct role in transcriptional repression but its binding sites overlap with insulator elements, indicating that L(3)mbt might also function as an insulator accessory factor (Richter et al., 2011; Van Bortle et al., 2014). Notably, L(3)mbt was purified in two non-enzymatic repressive chromatin complexes: the Drosophila RBF, E2F2, and Myb-interacting proteins (dREAM complex, also called Myb-Muv B) as well as the L(3)mbt-Interacting complex (LINT complex) (Lewis et al., 2004; Meier et al., 2012). DREAM is a large multi-subunit complex that controls gene expression throughout the cell cycle but also represses developmental genes. L(3)mbt associates at sub-stoichiometric levels with dREAM and is strictly found in its repressive forms (Georlette et al., 2007; Lewis et al., 2004). The LINT complex is composed of L(3)mbt, the novel transcriptional repressor Lint1 as well as the corepressor CoREST, and was shown to silence developmental genes in cultured cells (Meier et al., 2012). Interestingly, dREAM and LINT complexes repress overlapping sets of genes in somatic cells, including genes that are normally expressed in the germline. Despite extensive biochemical studies, we still know little about which chromatin complex mediates L(3)mbt’s role in tissue identity.

*Drosophila melanogaster* ovaries are composed of 16 to 20 egg assembly chains called ovarioles (Fig. 1A,B). At the tip of each ovariole a region called the germarium houses germline stem cells (GSCs), which divide asymmetrically to generate a new GSC and a differentiating daughter cell. The differentiating GSC daughter undergoes four rounds of mitosis with incomplete cytokinesis to form a sixteen-cell germline cyst in which sibling germ cells remain interconnected through cytoplasmic bridges called ring canals. GSCs are marked by a spectrin-containing spherical Endoplasmic Reticulum-derived vesicle known as spectrosome, which fuses into a branched fusome connecting the cells of the same cysts through the ring canals (Huynh, 2006). Only one of the cyst germ cells develops into an oocyte, while the other 15 cells become supportive, polyploid nurse cells. Somatic cells of the ovary play important roles in supporting oogenesis: they compose the GSC niche that promotes cyst divisions and differentiation, and the follicle cells enclose and individualize egg chambers facilitating oocyte-nurse cell development.

**Figure 1.**
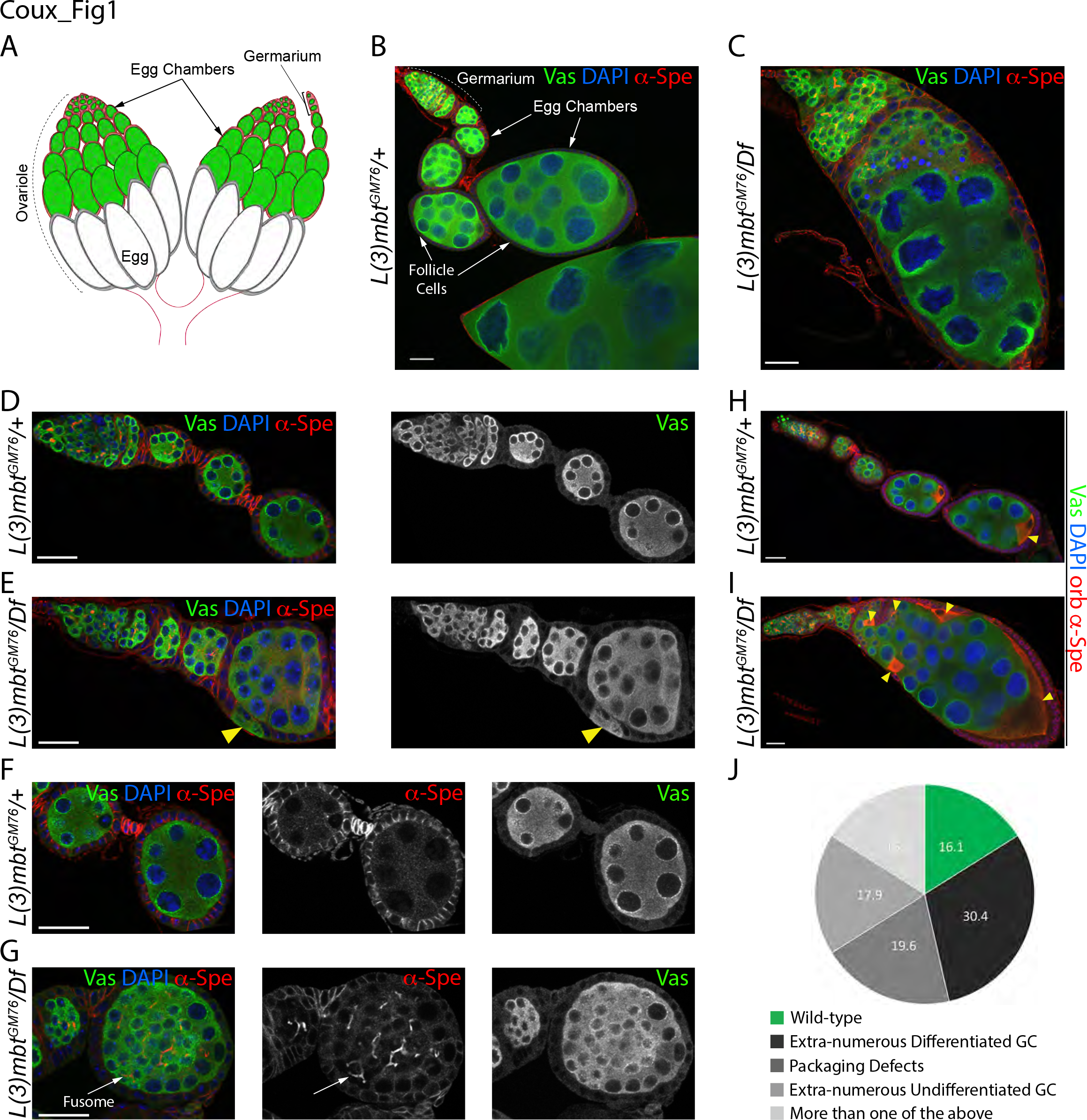
Developmental defects of *l(3)mbt* mutant ovaries. *(A)* Schematic representation of a wild-type ovary composed of ovarioles. *(B)-(G)* Confocal images of control and *l(3)mbt* mutant ovarioles stained for germ cells (Vasa, green), α-Spectrin (red), and DAPI (blue) for DNA. All ovarioles images are displayed with anterior oriented to the top-left corner. *(B)* Heterozygous control ovariole. *(C)* Representative *l(3)mbt* mutant ovariole with extra-numerous undifferentiated and differentiated germ cells surrounded by follicle cells. *(D)* Tip of wild-type ovariole with germarium and early egg chambers. *(E)* Mutant ovariole with germline packaging defects showing Vasa-expressing germ cells intercalated between follicle cells (yellow arrow). *(F)* Wild-type stage 3 and 4 egg chambers. Germ cells within egg chamber are no longer connected by fusomes. *(G)* Similarly staged mutant egg chamber filled with fusome-containing undifferentiated germ cells (arrow). *(H,I)* Confocal images of control and mutant ovarioles stained for Vasa (green), Orb (oocyte marker), α-Spectrin (red) and DAPI (blue). *(I)* In control ovarioles, Orb is restricted to the developing oocyte at the posterior of egg chambers. *(J) l(3)mbt* mutant ovariole with an egg chamber containing more than 16 germ cells and multiple oocytes, as revealed by Orb staining. *(J)* Quantification of phenotypes observed in *L(3)mbt* mutant ovarioles as illustrated in (*C, E and G*) Scale bars, 25 μm.

To understand how L(3)mbt secures tissue identity, we combined genetic and genomic approaches to characterize the functions of L(3)mbt in *Drosophila melanogaster* ovarian development. We find that L(3)mbt affects gene expression in a tissue-specific manner. In somatic cells of the ovary, L(3)mbt represses germline genes, whereas in the female germline it controls genes normally expressed in the testis and the nervous system. Mutant ovarian tissues continue to express signatures of the tissue of origin, indicating that loss of L(3)mbt does not induce transdifferentiation. Remarkably, we show that ectopic expression of a single gene in the somatic ovarian cells, the translational repressor and key regulator of germline fate *nanos*, is largely responsible for aberrant development. Using a genetic approach, we find that in the ovary L(3)mbt function requires its cofactor Lint1 but is independent of the dREAM complex. Together, our experiments provide insight into the role of L(3)mbt in securing tissue identity by repressing expression signatures characteristic of other tissues that are incompatible with normal development.

## Results

### Sterility in *l(3)mbt* mutant females is associated with aberrant ovarian development

L(3)mbt was previously shown to be required for the development of the nervous system as *l(3)mbt* mutant flies grown at restrictive temperatures (29°C) develop malignant brain tumors and die at larval stages (Gateff et al., 1993; Janic et al., 2010; Richter et al., 2011). We observed that, when grown at lower temperature (18°C or 25°C), null *l(3)mbt* mutant females were viable but fully sterile, indicating that L(3)mbt is critically required for germline development. At the macroscopic level, *l(3)mbt* mutant ovaries were atrophied and adult females did not lay eggs. To characterize the ovarian phenotype in detail, we used antibodies against the germline marker Vasa and α-Spectrin (α-Spe), which labels the membranes of somatic cells and spectrosomes/fusomes. Similar to the wild type, *l(3)mbt* mutant germaria contained GSCs adjacent to the somatic niche (Fig. S1 A-B). However, mutant ovarioles contained fewer individualized egg chambers (1.35 egg chambers/ovariole in average vs. 6 in WT, p<10^−4^), which were highly abnormal and displayed several defects (Fig. 1C, E, G, quantified in J). Such defects included extra-numerous germ / nurse cells within egg chambers (Fig. 1C; “extra-numerous differentiated germ cells” in Fig. 1J) as well as egg chambers in which somatic follicle cells fail to fully enclose germ cells (Fig. 1E, arrowhead; “packaging defects” in Fig 1J). In the wild type, fusomes degenerate as cysts proceed past the 16 cell stage and are enveloped by somatic follicle cells (Huynh and St Johnston, 2004). In *l(3)mbt* mutants, however, we observed accumulation of undifferentiated germ cells, which were marked by branched fusomes and enclosed by follicle cells (Fig. 1G, arrow; “extra-numerous undifferentiated germ cells” in Fig 1J).

Another hallmark of the 16-cell wild-type cyst is the specification of a single oocyte, while the remaining fifteen cells develop into polypoid feeder cells, the nurse cells. Using the RNA binding protein oo18 RNA-binding protein (Orb, Christerson and McKearin, 1994) as a marker for the future oocyte (figure 1H, arrowhead), we observed that most mutant egg chambers had multiple Orb positive cells (Fig. 1I, arrowheads). Wild-type oocytes are connected to the adjacent nurse cells by four ring canals, as a product of four divisions (Huynh and St Johnston 2004). We determined whether the additional cells were bona fide oocytes by counting the associated ring canals (stained by F-Actin, Fig. S1C and D, yellow arrows and dotted circles) and found that ectopic Orb-expressing cells contained four or more ring canals, while egg chambers contained multiple oocytes and extra-numerous germ cells. Thus, *l(3)mbt* loss causes egg chamber fusions and possibly, additional rounds of cyst division. In rare cases (2%), we observed multiple germaria connected to the same aberrant egg chamber (Fig. S1 E,F), suggesting that ovarioles were fused during ovary morphogenesis. Taken together, our results indicate that in addition to its previously reported, conditional requirement in the brain, L(3)mbt has a critical, temperature-independent role required for ovarian morphogenesis and differentiation.

### L(3)mbt functions in ovarian somatic cells to safeguard ovary development

Previous experiments had shown that loss of *l(3)mbt* specifically in germ cells caused developmental defects in the resulting embryos, but allowed oocytes to mature (Yohn et al., 2003). Thus, we wondered whether the gross abnormalities of mutant ovaries were indicative of a role for L(3)mbt in somatic cells of the ovary. To determine the tissue-specific requirement of L(3)mbt, we generated homozygous *l(3)mbt*^*GM76*^ mutant clones in the ovarian somatic cells by using the FRT-FLP system (Harrison and Perrimon, 1993) under the transcriptional control of the *Ptc*-*Gal4* or *c587*-*Gal4* drivers, which drive expression in the somatic cells of the germarium (Hinz et al., 1994; Zhu and Xie, 2003). Interestingly, loss of L(3)mbt in a subset of somatic cells perturbed germline development leading to egg chambers that contained extra-numerous germ cells (Fig. 2A) or multiple oocytes (Fig. 2B). To conclusively test for a role of L(3)mbt in somatic ovarian cells, we expressed an inducible *UAS*-*l(3)mbt::myc* transgene under the control of the tj-Gal4 driver and found that expression of L(3)mbt in the somatic cells of the ovary alone was sufficient to rescue the aberrant morphology of mutant ovaries, including numbers of oocytes and ring canal (Fig. 2C,D and Fig. S2A,B). These results demonstrate that L(3)mbt is required specifically in the somatic tissues of the ovary to support normal oogenesis.

**Figure 2.**
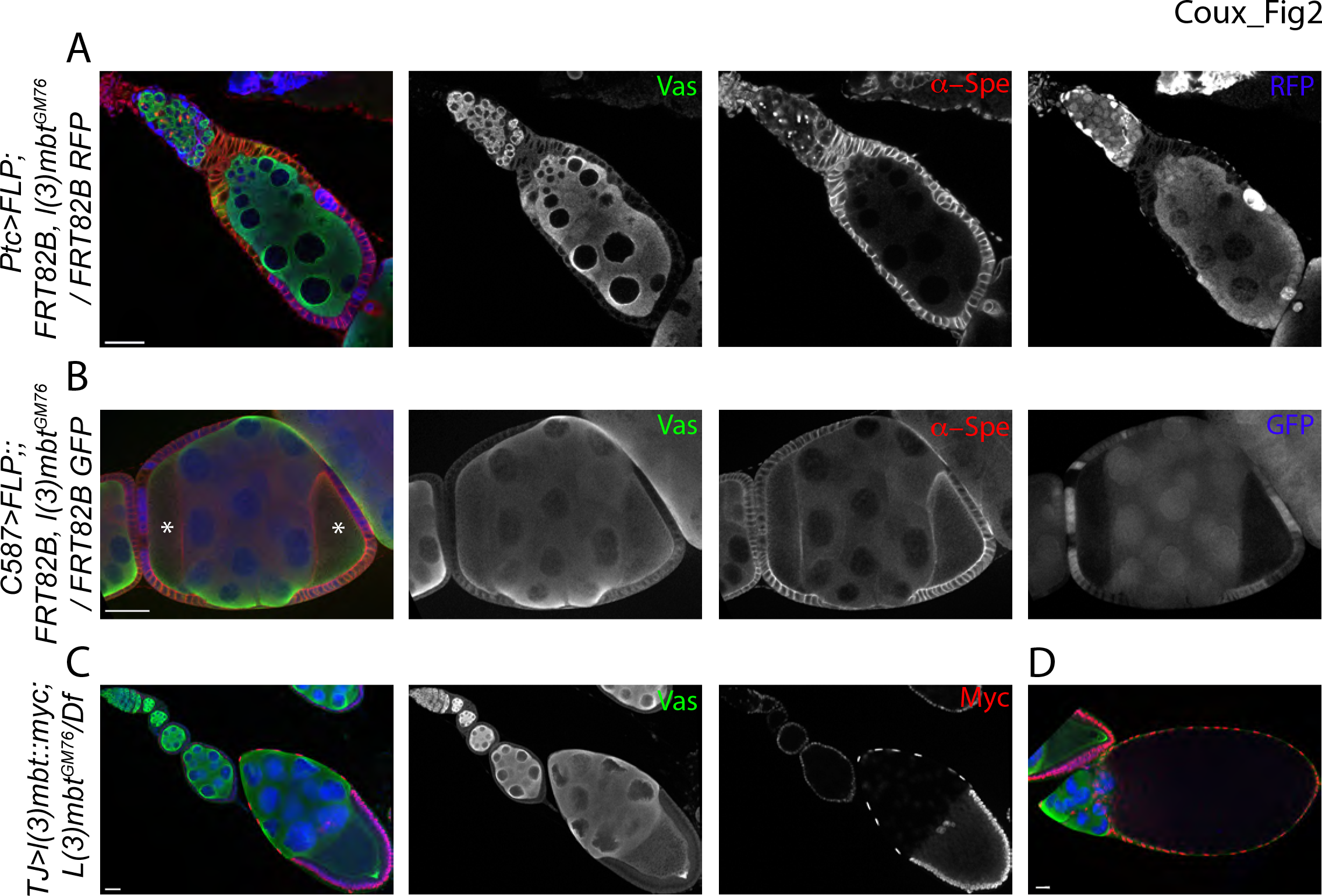
L(3)mbt functions in somatic cells for ovary development. *(A,B)* Confocal images of representative ovarioles with mutant *l(3)mbt* follicle cell clones marked by absence of RFP (A) or GFP (B) (blue), Vasa (green), α-Spectrin (red). Egg chambers surrounded by numerous *l(3)mbt* mutant follicle cells exhibits aberrant phenotypes. Oocytes are marked by asterisks in *(B)*. *(C,D)* Confocal images of mutant ovaries expressing *TJ*>*UAS-l(3)mbt::myc* in somatic cells stained for Vasa (green), Myc (red), and DAPI (blue). *(C)* Ovarian morphology and germ cell number are fully rescued by expression of a L(3)mbt wild-type transgene in somatic cells. *(D)* Rescued late stage oocyte. Scale bars, 25 μm.

### Mutant larval somatic cells are properly specified but intermingled cells fail to contact with germ cells

Ovarioles lacking L(3)mbt exhibit striking morphological defects. Most of the structures in the adult ovary are established and organized during the third instar larval (L3) stage (Gilboa, 2015), we thus examined ovaries from mid to late L3 larvae to investigate whether *l(3)mbt* mutation affects ovarian organogenesis. Germ cells and somatic cells associate during late embryonic stages and proliferate during most of larval development. Starting mid L3 stage, the somatic precursors differentiate into distinct populations of somatic cells (Gilboa and Lehmann, 2006; Li et al., 2003). In the apical compartment, post-mitotic terminal filament cells stack to form terminal filaments and associate with sheath cells (Godt and Laski, 1995). In the medial region, the Intermingled cells (IC) are closely associated with germ cells and are thought to give rise to the adult escort cells (Gilboa, 2015). We performed confocal imaging analysis by immunostaining with antibodies against the transcription factor Traffic Jam (TJ), which has important functions in specifying somatic gonadal cell types and labels the ICs (Li et al. 2003), Vasa and α-Spectrin. We observed that, like their wild-type counterparts, *l(3)mbt* mutant L3 ovaries have distinct apical compartments harboring terminal filaments, as well as a medial region containing germ cells and ICs (Fig. 3A,B). ICs are normally scattered throughout the germ cell population, however mutant ICs were excluded from the germ cell-containing region (Fig. 3B). Despite this aberrant behavior, *l(3)mbt* mutant ICs retained expression of Zfh1, a transcription factor essential for the somatic fate (Maimon et al., 2014, Fig. S3). Taken together, our results show that in *l(3)mbt* mutant L3 ovaries, markers for somatic cell fates are expressed but spatial organization is affected.

**Figure 3.**
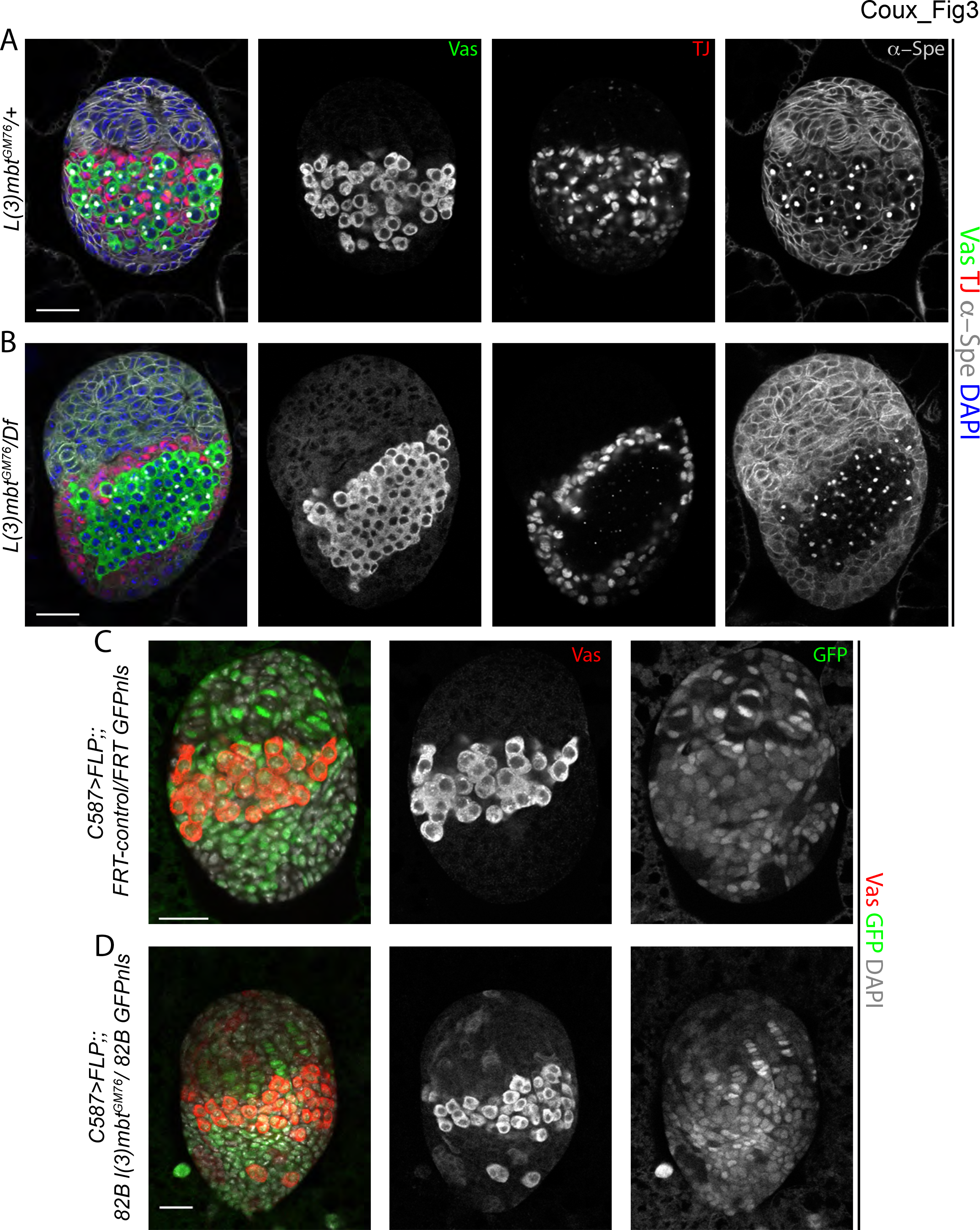
Larval somatic cells are properly specified but derepress Vasa. *(A)* Wild-type and *(B) l(3)mbt* mutant L3 ovaries stained for Vasa (green), TJ (red), α-spectrin (grey), and DAPI (blue). Scale bars represent 25*μ*m. *L(3)mbt* mutant ICs fail to migrate in between PGCs and are found around them. *(C,D)* Confocal images of L3 ovaries with wild-type *(C)* or *l(3)mbt* mutant clones *(D)* marked by the absence of GFP. stained for Vasa (red), GFP (green) and DAPI (grey). *l(3)mbt*^*GM76*^ mutant clones (labeled by the absence of GFP) in somatic cells express Vasa while wild-type clones do not.

### *L(3)mbt* mutant somatic larval cells ectopically express the germline marker *vasa*

Previous studies suggested that L(3)mbt loss results in derepression of germline genes in the larval brain and cultured somatic cells (Georlette et al., 2007; Janic et al., 2010; Meier et al., 2012; Sumiyoshi et al., 2016). Thus, we asked whether germline genes were ectopically expressed in mutant larval somatic cells. Indeed, we observed faint Vasa antibody signal in the somatic tissues of *l(3)mbt* mutant ovaries, especially in the apical compartment (Fig. 3B and S3). To further demomstrate these initial observations, we induced *l(3)mbt* mutant clones using the c587-Gal4 driver to express FLP recombinase specifically in the somatic tissues of the larval ovary (Zhu and Xie, 2003). Induction of wild-type control clones (labeled by the absence of GFP) in somatic cells of the basal or apical compartment did not cause Vasa expression (Fig. 3C). In contrast, *l(3)mbt*^*GM76*^ homozygous clones of somatic cells exhibited Vasa staining (Fig. 3D). From this, we conclude that L(3)mbt represses the germ cell marker *vasa* in the somatic cells of the larval ovary.

### *L(3)mbt* mutant somatic ovarian cells simultaneously express somatic gonad and germline-specific genes

To gain a genome-wide view of gene expression changes induced by loss of L(3)mbt in adult somatic ovarian cells *in vivo*, we performed RNA-sequencing (RNA-seq) analysis. To distinguish between germline and somatic ovarian tissues, we took advantage of *tud* maternal mutations (*tud*^*M*^), which give rise to progeny that lack germ cells and develop into adults devoid of germline (Arkov et al., 2006; Smendziuk et al., 2015; see File S1 and Materials and Methods). Comparisons between *tud*^*M*^; *l(3)mbt*^*GM76*^/*l(3)mbt*^*Df*^ and *tud*^*M*^; *l(3)mbt*^*GM76*^/+ adult ovaries identified 600 genes differentially expressed (*adjusted p*-*value*<0.05) in the somatic cells of the ovary. Of these, 459 were upregulated and 141 downregulated in mutant tissues (Table S1). 44 upregulated genes were shared with the 101 MBTS genes (Janic et al., 2010), and 115 out of the 681 genes found upregulated in *Δ*-*l(3)mbt* Ovarian Somatic Cells (OSC; Sumiyoshi et al., 2016). These derepressed genes include piRNA pathway components and germline-specific genes such as *nos*, *Pxt*, *vas*, *aub*, *tej*, *krimp*, *AGO3*, and *CG9925* (Fig. 4A, Fig. S4A). The effect of *l(3)mbt* mutation on gene expression was more pronounced for repressed genes, while genes normally expressed in the soma showed only low fold-changes in the mutant (124/141 had a Log_2_ fold-change between −0.3 and −1). To investigate whether mutant somatic cells retained their somatic identity, we performed immuno-stainings against the key transcription factors Traffic Jam (TJ) and Zfh1, which are essential for gonad development and exclusively expressed in somatic cells (Leatherman and Dinardo, 2008; Li et al., 2003). Despite the gross morphological abnormalities, we observed TJ or Zfh1 expressing cells surrounding germ cells in mutant ovaries (Fig. 4B,C and Fig. S4B,C). Further, F-Actin staining showed that mutant somatic cells retain the columnar morphology and epithelial characteristics of wild-type follicle cells (Fig. S1D and S2A). However, individual TJ-positive cells occasionally expressed the germline marker Vasa (Fig. 4C, arrows). As TJ/Zfh1 and Vasa expressions are mutually exclusive in wild-type ovaries, our results indicate that *l(3)mbt* mutant somatic cells retain somatic and epithelial/follicular characteristics while ectopically expressing hallmark genes of germline fate.

**Figure 4.**
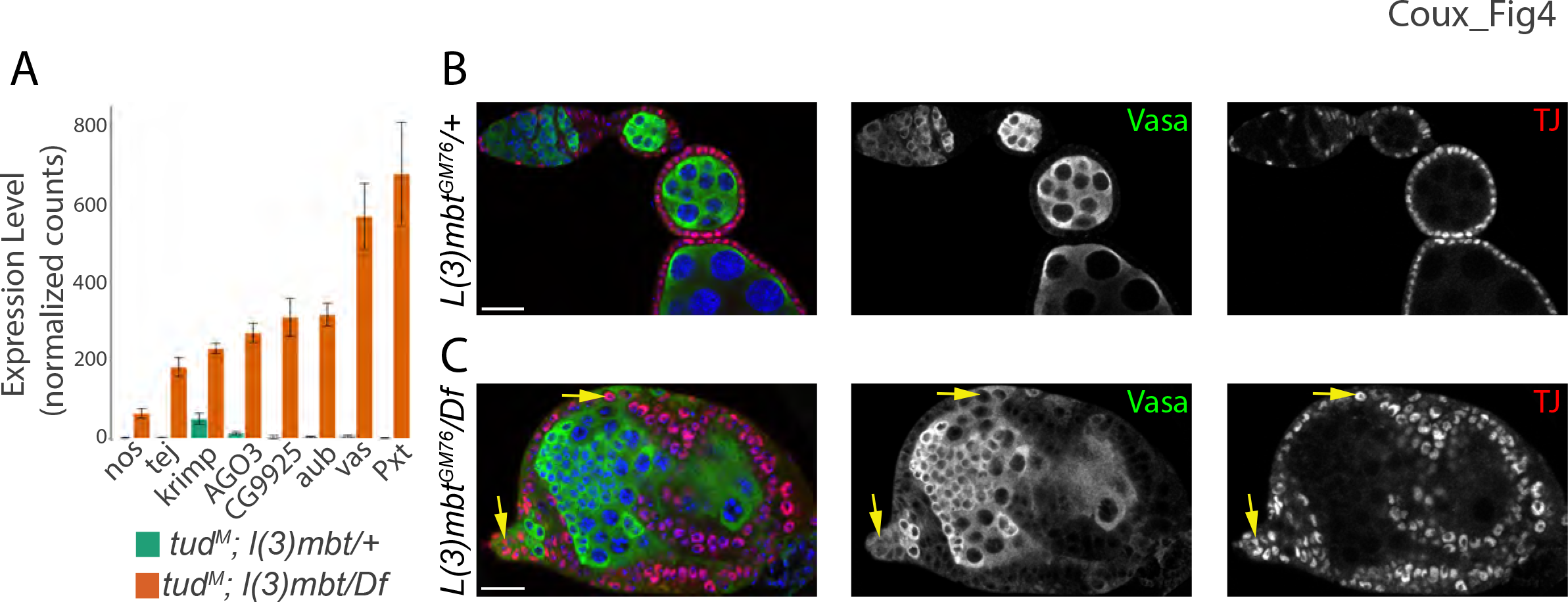
*l(3)mbt* mutant somatic cells are properly specified but ectopically express germline genes. *(A)* Expression level of the germline-specific genes *nos*, *tej*, *krimp*, *AGO3*, *CG9925*, *aub*, *vas* and *Pxt* in *tud*^*M*^ ovaries heterozygous and homozygous mutant for *l(3)mbt*, as measured by RNA-seq analysis (expressed in normalized counts). *(B*,*C)*focal images of control and *l(3)mbt* mutant ovarioles stained for Vasa (green), Traffic-Jam (TJ; red) and DAPI (blue). TJ is expressed in all somatic cells of the adult ovary. *(C)* Some TJ-positive somatic cells express the germline marker Vasa (yellow arrows). Scale bars, 25 μm.

### Expression of Nos is necessary and sufficient to cause ovarian defects

Aberrant growth of *l(3)mbt* brain tumors was shown to rely on the ectopic expression of *nos*, *aub*, and *vasa* (Janic et al., 2010). We find that each of these genes is indeed upregulated in compromised somatic ovarian cells lacking L(3)mbt (Fig. 4A and S4A). We therefore asked whether their misexpression contributed to the ovarian defects observed of *l(3)mbt* mutants by generating double mutant animals. Due to lethality, we were unable to assess ovarian phenotypes in *vas*, *l(3)mbt* mutants. However we found that *aub*, *l(3)mbt* double mutant ovaries were phenotypically similar to *l(3)mbt* single mutant, containing many apoptotic cells and aberrant egg chambers with more than 16 germ cells (Fig. S5A). In contrast, *nos*^*BN*^/*nos*^*L7*^ mutations dramatically suppressed the *l(3)mbt* ovarian defects: *nos*, *l(3)mbt* double mutant ovaries contained late stages egg chambers with ovarioles phenotypically indistinguishable from wild type and 85% of double mutant egg chambers contained 16 germ cells and only one oocyte (Fig. 5C-E, n=88). Consistently, depletion of Nanos’ co-factor Pumilio in *l(3)mbt* mutant ovaries also suppressed the mutant phenotypes, with 81% of *pum*^680^, *l(3)mbt* double mutant ovarioles resembling wild-type morphology (Fig. 5F-H, n=181). These data suggest that Nanos is a critical factor leading to *l(3)mbt* mutant ovarian phenotypes. To test whether *nos* misexpression in somatic ovarian cells is sufficient to cause *l(3)mbt*-like ovarian defects, we ectopically expressed *nos* in the somatic cells of the ovary using the tj-Gal4 driver. *Nos* misexpression during larval stages caused lethality, we therefore restricted *nos* expression in the soma to adult stages using the Gal80^ts^ system (McGuire et al., 2004). *Nos* somatic expression perturbed ovarian morphology resulting in defects of the follicle epithelium and poorly individualized egg chambers (Fig. S5B). While this phenotype resembles the *l(3)mbt* phenotype, it does not fully recapitulate it, possibly because *nanos* was only misexpressed in adult tissues likely at higher levels compared to *l(3)mbt* mutants. In contrast, ectopic expression of *aub* or *vas* in somatic ovarian cells did not yield a morphologically significant phenotype (Fig. S5C, D). Together, these results suggest that ectopic expression of *nos*, but not *aub or vas*, is necessary and sufficient to cause aberrant somatic ovarian development.

**Figure 5.**
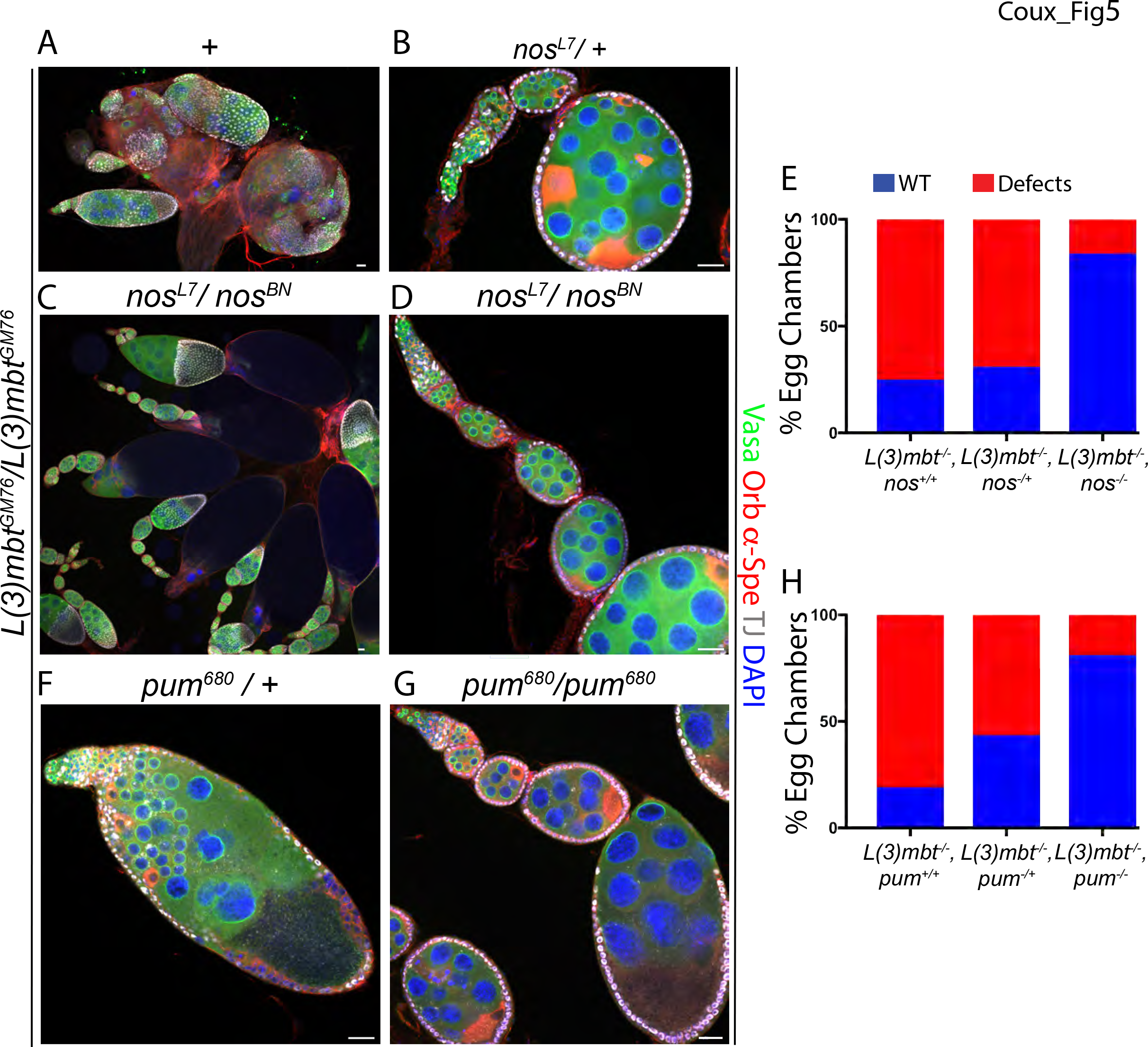
Nanos and its cofactor Pumilio mediate developmental phenotypes in *l(3)mbt* ovaries. *(A-D,F-G)* Confocal images of ovaries stained for Vasa (green), Orb and α-Spectrin (red), TJ (grey), and DAPI (blue). *(A-D)* Representative confocal images of (*A*) *l(3)mbt*^*GM76*^, (*B*) *l(3)mbt*^**GM76**^, *nos*^*L7*^/ *l(3)mbt*^*GM76*^, +, *and (C-D)l(3)mbt*^*GM76*^, *nos*^*L7*^ / *l(3)mbt*^*GM76*^, *nos*^*BN*^ double mutant ovarioles. *(A,C)* Low magnification and *(B,D)* medium magnification images. *(E)* Quantification of phenotypes observed in the genotypes described in *(A-D)*. *(F-G)* Confocal images of (*F*) *l(3)mbt*^*GM76*^, *pum*^*680*^ / *l(3)mbt*^*GM76*^, + and (*G*) *l(3)mbt*^*GM76*^, *pum*^680^ homozygous ovarioles. *(H)* Quantification of phenotypes observed in the genotypes described in *(F-G)*. Scale bars, 25 μm.

### L(3)mbt functions through the LINT complex to secure ovarian development

L(3)mbt has been associated with two chromatin complexes that repress developmental and germline genes: the dREAM and LINT complexes (Fig. 6A-B; Georlette et al., 2007; Meier et al., 2012). To determine the function of these complexes in ovarian development, we depleted somatic cells from E2f2 and Mip120, two core repressors of the dREAM complex. Mutant somatic clones for *E2f2* (Fig. 6C) or ovaries deficient for *mip120* (Fig. 6E) did not result in phenotypic aberrations reminiscent of those observed in *l(3)mbt* mutants. We also asked whether *mip120* mutation affected L(3)mbt nuclear localization in the ovary, as previously described in salivary glands (Blanchard et al., 2014). Using the TdTomato::L(3)mbt fusion protein we observed L(3)mbt nuclear localization comparable to wild type in *mip120*^*67.9A.9*^ mutants (Fig. S6B). Together, these results suggest that L(3)mbt critical function in somatic ovarian cells is independent of the dREAM complex.

**Figure 6.**
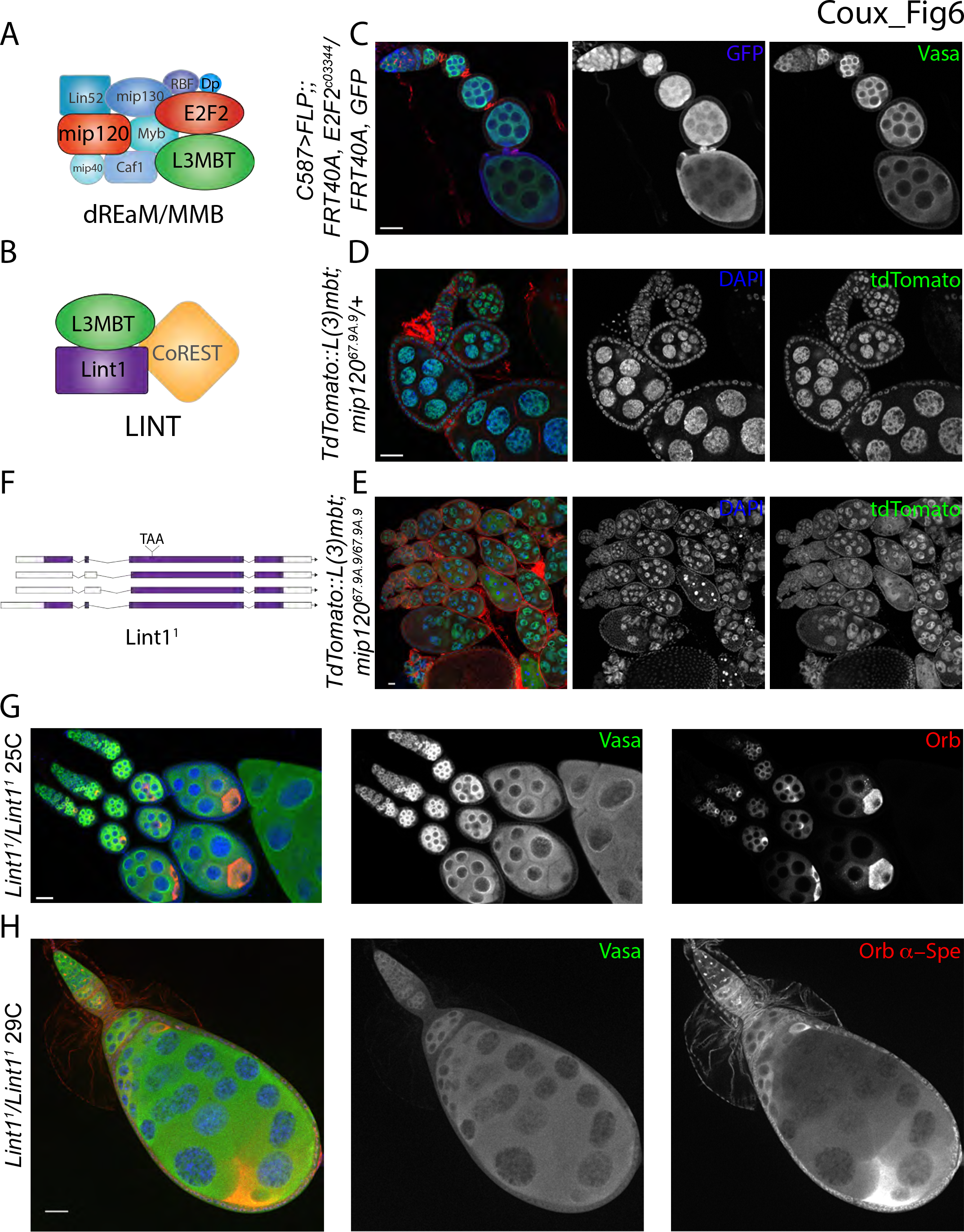
LINT complex mutants have ovarian defects similar to *l(3)mbt*. *(A-B)* Schematic representation of (*A*) the dREAM/MMB and (*B*) LINT complexes. (*C*) Confocal image of ovariole with *E2f2*^*c03344*^ mutant clones marked by absence of GFP (blue), Vasa (green), α-Spectrin (red). *(D-E)* Confocal images of control and *mip120*^*67.A9.9*^ mutant ovaries expressing the tdTomato::L(3)mbt fusion and stained for α-Spectrin (red), tdTomato (green), and DAPI (blue). *(D) mip120*^67.A9.9^ heterozygous control ovarioles. *(E)* Low magnification confocal image of homozygous mutant *mip120*^67.A9.9^ ovarioles accumulating stage 7-9 egg chambers that eventually undergo apoptosis. *(F)* Schematic representation of the *Lint1*^*1*^ allele. *(G-H)* Confocal images of Lint1 mutant ovaries stained for Vasa (green), Orb (red), and DAPI (blue), α-Spectrin (red) in *(H)*. *Lint1*^*1*^ mutants grown at (G) At 25°C, few mutant egg chambers contain two Orb-positive cells or mis-positioned oocytes. (H) At 29°C mutant egg chambers can contain more than 16 germ cells and multiple oocytes similar to defects observed in *l(3)mbt* mutants. Scale bars, 25 μm.

Next we examined the role of the LINT complex as a mediator of L(3)mbt function. Since mutations in *Lint1* had not been identified, we generated CRISPR-induced *Lint1* alleles (Gratz et al., 2013; Gratz et al., 2014). *Lint1*^*1*^ deletes two cytosines (350 and 351), creating a premature stop codon at position 66/540 (Lint1-C) or 128/601 (Lint1-A, Fig. 6F). Homozygous mutant flies were viable and fertile at 25°C, and their ovaries developed normally although 2% of egg chambers contained misplaced or extra-numerous oocytes (Fig. 6G). However, when grown at 29°C, *lint1*^*1*^ females were fully sterile, laying eggs that failed to hatch, and 15% of their ovarioles developed aberrantly and exhibited *l(3)mbt* mutant phenotype (Fig. 6H). Furthermore, depletion of one *l(3)mbt* copy rendered *lint1*^*1*^ homozygous females (*lint1*^*1*^/ *lint1*^*1*^*;;l(3)mbt*^*GM76*^/+) fully sterile at 25°C, confirming the genetic interaction. We conclude that L(3)mbt’s function in the ovary is mediated by its cofactor Lint1 and the LINT complex.

### L(3)mbt is autonomously required in the germline for egg chamber survival and represses neuronal and testis-specific genes in the female germline

In addition to its role in the development of the somatic cells of the Drosophila ovary, L(3)mbt also has a maternal, germline-autonomous role that supports nuclear divisions during early embryogenesis (Yohn et al., 2003). Similarly, we observed that embryos laid by mutant females, which somatically expressed the complementing *l(3)mbt::myc* transgene, failed to hatch. Further, while most egg chambers appeared morphologically normal in somatically-complemented *l(3)mbt* mutant ovaries, we noticed that 69% of ovarioles contained apoptotic egg chambers (Fig. 7A,B). This suggests that in addition to its previously identified maternal effect function for early embryonic development, L(3)mbt is autonomously required in the germline for egg chamber development.

**Figure 7.**
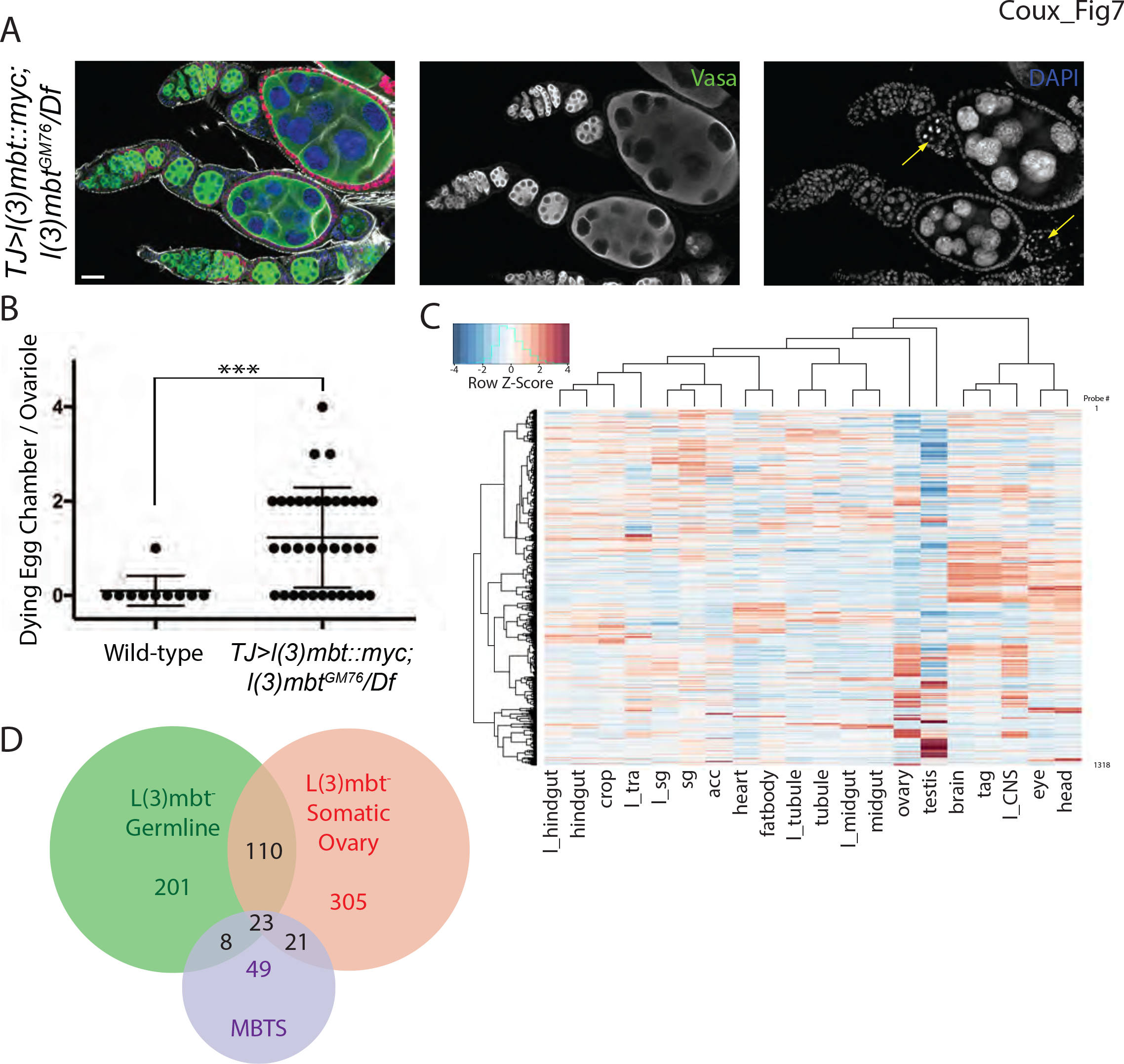
L(3)mbt represses neuronal and testis-specific genes in the germline. *(A)* Confocal image of *l(3)mbt* mutant ovarioles expressing the *l(3)mbt::myc* transgene in somatic cells, stained for Vasa (green), Myc (red), F-Actin (grey), and DAPI (blue), Scale bars, 25 μm. *L(3)mbt* mutant egg chambers surrounded by somatic cells expressing the L(3)mbt::myc fusion undergo cell death (yellow arrows). *(B)* Quantification of egg chambers undergoing cell death. *(C)* Hierarchical clustering of tissue expression profile of genes repressed by L(3)mbt in the female germline. Gene expression per tissue (normalized to fly average) is shown as a Z-Score heatmap. *(D)* Venn Diagram showing genes upregulated in *l(3)mbt* mutant ovarian soma (red), female germline (green), and larval brain tumors (Janic et al., 2010) (MBTS, purple). Most derepressed genes are tissue specific.

Our data highlight a role for L(3)mbt in suppressing germline specific genes in somatic tissues of the ovary. However, our experiments also uncovered a germline autonomous requirement for egg chamber development. To gain a genome-wide view of the changes in gene expression specifically induced by loss of L(3)mbt in the germline, we performed RNA-seq analysis on embryos laid by *l(3)mbt* mutant or heterozygous mothers expressing the *L(3)mbt::myc* fusion in the somatic ovary. We used early embryos prior to activation of the zygotic genome as the early embryonic RNA pool is exclusively composed of maternally provided transcripts and thereby reflects the germline expression profile during oogenesis (Edgar and Schubiger, 1986). Our analysis identified 878 differentially expressed genes (*adjusted p*-*value*<0.05), of which 342 were upregulated and 536 downregulated in embryos laid by mutant females (Table S2). Most upregulated genes were uncharacterized and not enriched for specific Gene Ontology terms. Thus, to better characterize the group of genes upregulated in the *l(3)mbt* mutant germline, we performed two-way hierarchical clustering to identify any tissue specific expression signatures. Two major groups were readily identified: one composed of genes highly expressed in neuronal tissues (*brain*, *thoracicoabdominal ganglion (tag)*, *larval CNS*, *eye*, and *head*; Fig. 7C) and another comprising of testis-specific genes. As in mutant somatic ovaries, two-thirds (365/536) of the downregulated genes had their expression level reduced by less than two-fold compared to the heterozygous controls (Fig. S7). We observed only a limited overlap between the genes derepressed in *l(3)mbt* mutant somatic and germline ovarian tissues, with two thirds of genes being specifically upregulated in one of the two tissues (Fig. 7D). Taken together, these results indicate that L(3)mbt function is not restricted to repressing the germline program in somatic tissues, but that L(3)mbt regulates distinct sets of genes, in a tissue-specific manner.

## Discussion

By combining developmental and molecular analysis, we show that *l(3)mbt* mutant ovaries develop aberrantly. L(3)mbt depletion does not result in complete transdifferentiation but causes simultaneous expression of original cell signatures and ectopic expression of markers of other cell fates. We hypothesize that this conflict between co-existing cell identities causes the observed aberrant tissue morphogenesis. Direct support for this idea is provided by the role of the translational repressor and germline gene *nanos*, which derepression in somatic ovarian cells causes aberrant growth. Molecularly, we demonstrate that L(3)mbt functions through the LINT complex in the somatic ovary. Finally we show that L(3)mbt-mediated regulation of gene expreeion is not limited to repression of germline specific genes in somatic tissues is but tissue dependent. We propose that L(3)mbt functions, through LINT, as a guardian of cell identity by preventing the simultaneous expression of genes sets incompatible with such identity.

Our experiments demonstrate that ectopic expression of *nanos* is necessary and sufficient to induce aberrant development of *l(3)mbt* mutant ovaries. These defects are likely not due to a direct interference at the transcriptional level but are rather caused by Nos’ function as a translational repressor. In support, we find that Pumilio, the sequence-specific translational repressor and co-factor of Nos, is also essential for the *l(3)mbt* ovarian phenotype. Nos was recently shown to modulate Pum RNA-binding and target-specificity in somatic S2 cells (Weidmann et al., 2016). Since Pum is ubiquitously expressed, we propose that ectopic Nos stabilizes Pum binding at target mRNAs essential for somatic functions. Interestingly, ectopic expression of NANOS1 was found to be required for growth of human *pRb* deficient tumor cells. In this case, NANOS1 and PUM repress p53 translation allowing cells to bypass apoptosis (Miles et al., 2014). Thus, ectopic Nos-Pum complexes may alter tissue maintenance at the post-transcriptional level in other systems as well. As Nos has been found to repress somatic genes in germ cells of multiple organisms (Hayashi et al., 2004, Lee et al., 2017), we would expect to find key regulators of somatic fate among mRNAs aberrantly targeted by Nos-Pum in somatic Drosophila tissues.

In contrast to the widespread effects of *nos* derepression observed in multiple somatic tissues upon loss of L(3)mbt, ectopic expression of additional genes may define the exact phenotypic consequences, which depend on tissue type. For example, piRNA pathway genes are ectopically expressed in *l(3)mbt* larval brain tumors and somatic ovarian cells, however, depleting them ameliorates the brain tumor but not the ovarian phenotype. This difference may be explained by the fact that the somatic ovary uses core components of the piRNA pathway to regulate transposable elements (Handler et al., 2013). Similarly, we did not observe derepression of Hippo target genes in ovarian tissues. Consistent with our finding that in *l(3)mbt* mutants new and original tissue identities are co-expressed, these results suggest that the phenotypic consequences of *l(3)mbt* mutation depend on the context of the original tissue identity.

Our results demonstrate that L(3)mbt function in the ovary is independent of the dREAM complex. The dREAM complex has a well-established role in cell-cycle regulation (Sadasivam and Decaprio 2013). Indeed, Mip120, a core dREAM component, was recently found to be required for decondensation of nurse cell nuclei (Cheng et al., 2017) and E2F2 is required for endo-replication of follicle cells (Cayirlioglu et al., 2001). We did not observe a role for *l(3)mbt* in the regulation of nurse and follicle cell endo-replication. Instead, our data support the hypothesis that in the ovary, L(3)mbt functions predominantly through the LINT complex and that this complex can be functionally separated from of the dREAM complex. Considering the moderate phenotype of *lint1*^*1*^ mutants and that gene function is apparently dispensable at 25°C, we speculate that L(3)mbt exerts most of the repressive activity of this new complex, possibly with additional, yet unidentified interactors and that Lint1 has an accessory role.

Loss of L(3)mbt causes the ectopic expression of a number of genes including cell identify regulators that interfere with original cellular function and affect tissue development. In contrast to a previously suggested soma-to-germline transformation, our results favor the hypothesis that *l(3)mbt* mutation imbalances tissue-homeostasis whereby normally mutually exclusive lineage determinants become simultaneously expressed. In support, L(3)mbt depletion in neuronal, somatic ovarian, and germ cells does not lead to loss of original tissue-specific markers, but genes characteristic of other lineages are derepressed (Richter et al. 2011, this study). Moreover, our results suggest that the role of L(3)mbt is not solely restricted to prevent ectopic expression of germline genes, but instead L(3)mbt represses distinct, broader sets of genes in a tissue-specific manner. Therefore, we propose that L(3)mbt secures tissue identity by stabilizing gene expression profiles established during differentiation.

---

**Supplemental figure 1.**
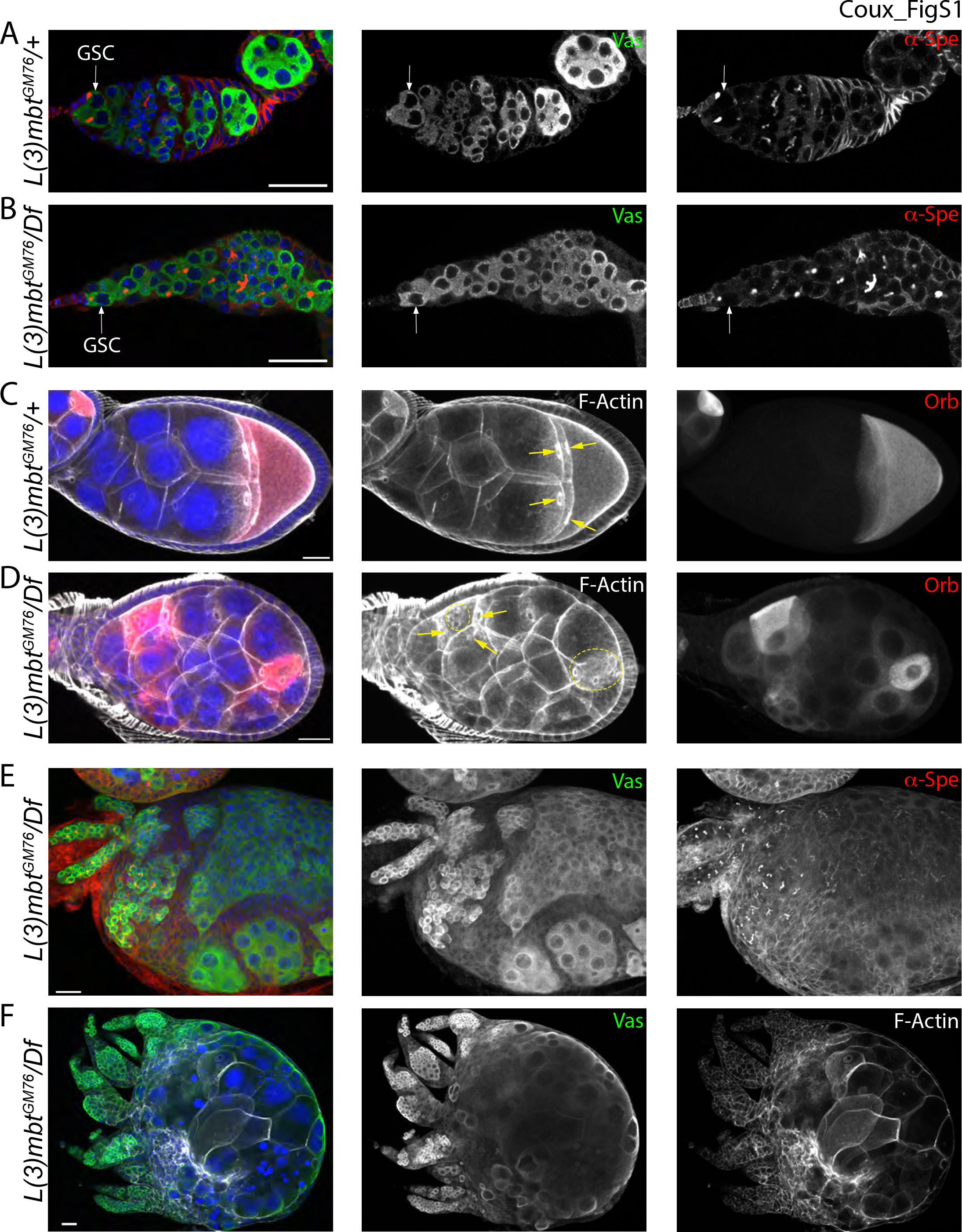
L(3)mbt loss-of-function leads to egg chamber and ovariole fusions. *(A-B)* Confocal images of wild-type *(A)* and mutant *(B)* germaria, stained for Vasa (green), α-Spectrin (red) and DAPI (blue). Wild-type and mutant germaria contain Germline Stem Cells marked by punctuated fusomes (α-Spectrin), in contact with the somatic niche. *(C-D)* Confocal projections of *(C)* wild-type and *(D)* mutant ovaries stained for F-Actin (grey), Orb (red), and DAPI (blue). Anterior is oriented to the left. *(C)* a wild-type oocyte is connected to nurse cells by four ring canals (yellow arrows). *(D) L(3)mbt* mutant egg chamber containing multiple oocytes with four or more ring canals (arrows and dotted circles). *(E)* Confocal image of *l(3)mbt* mutant ovary stained for Vasa (green), α-Spectrin (red) and DAPI (blue). Three germaria (top left) fused into an aberrant ovariole with intermingled differentiated and undifferentiated germ cells. *(E) l(3)mbt* mutant ovary with multiple germaria connected to the same giant egg chamber. Vasa (green), F-Actin (grey) and DAPI (blue). Scale bars, 25 μm.

**Supplemental figure 2.**
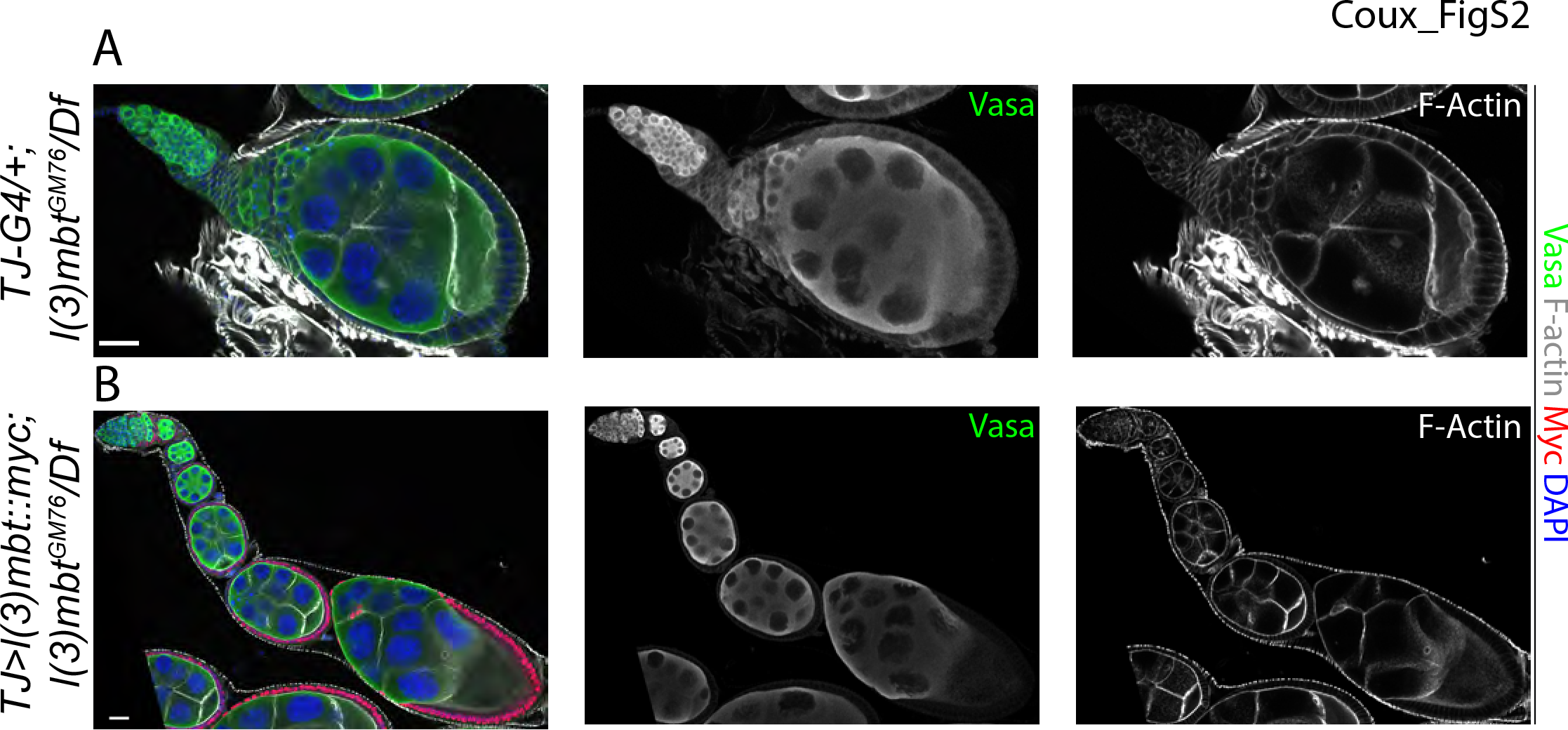
L(3)mbt somatic expression rescues *l(3)mbt* mutant ovarian morphology. Confocal images of ovarioles from (*A*) *l(3)mbt* mutant control and (*B*) *l(3)mbt* mutant expressing the *l(3)mbt::myc* transgene in somatic cells, stained for Vasa (green), Myc (red), F-Actin (grey), and DAPI (blue). Complemented ovarioles show wild-type morphology, including proper oocyte specification and ring canals numbers. Scale bars, 25 μm.

**Supplemental figure 3.**
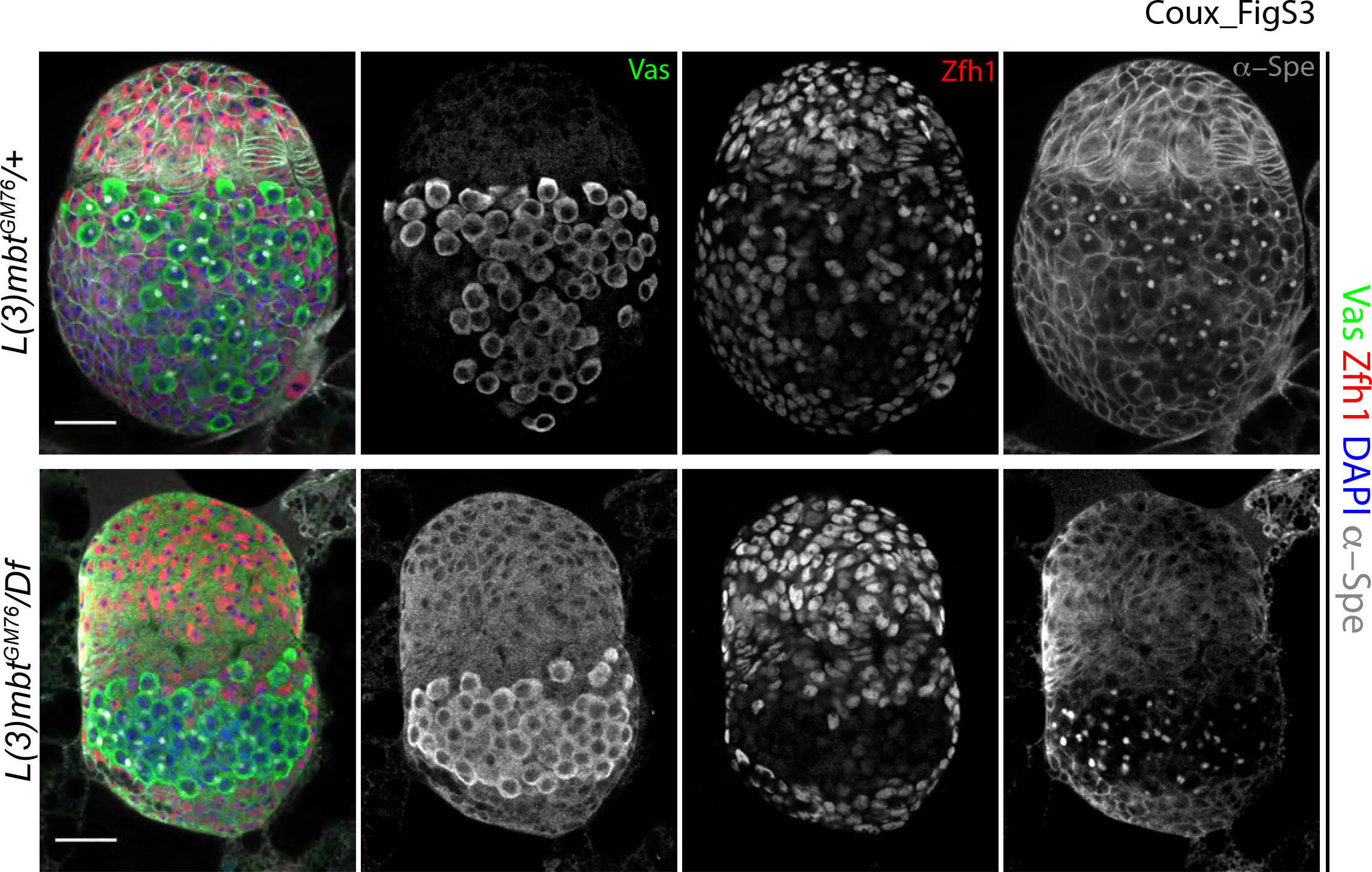
*L(3)mbt* mutant somatic larval cells have normal Zfh1 expression. Wild-type (top panel) and *l(3)mbt* mutant (bottom). L3 ovaries stained for vasa (green), α-spectrin (grey), Zfh1 (red) and DAPI (blue). Scale bars represent 25*μ*m.

**Supplemental figure 4.**
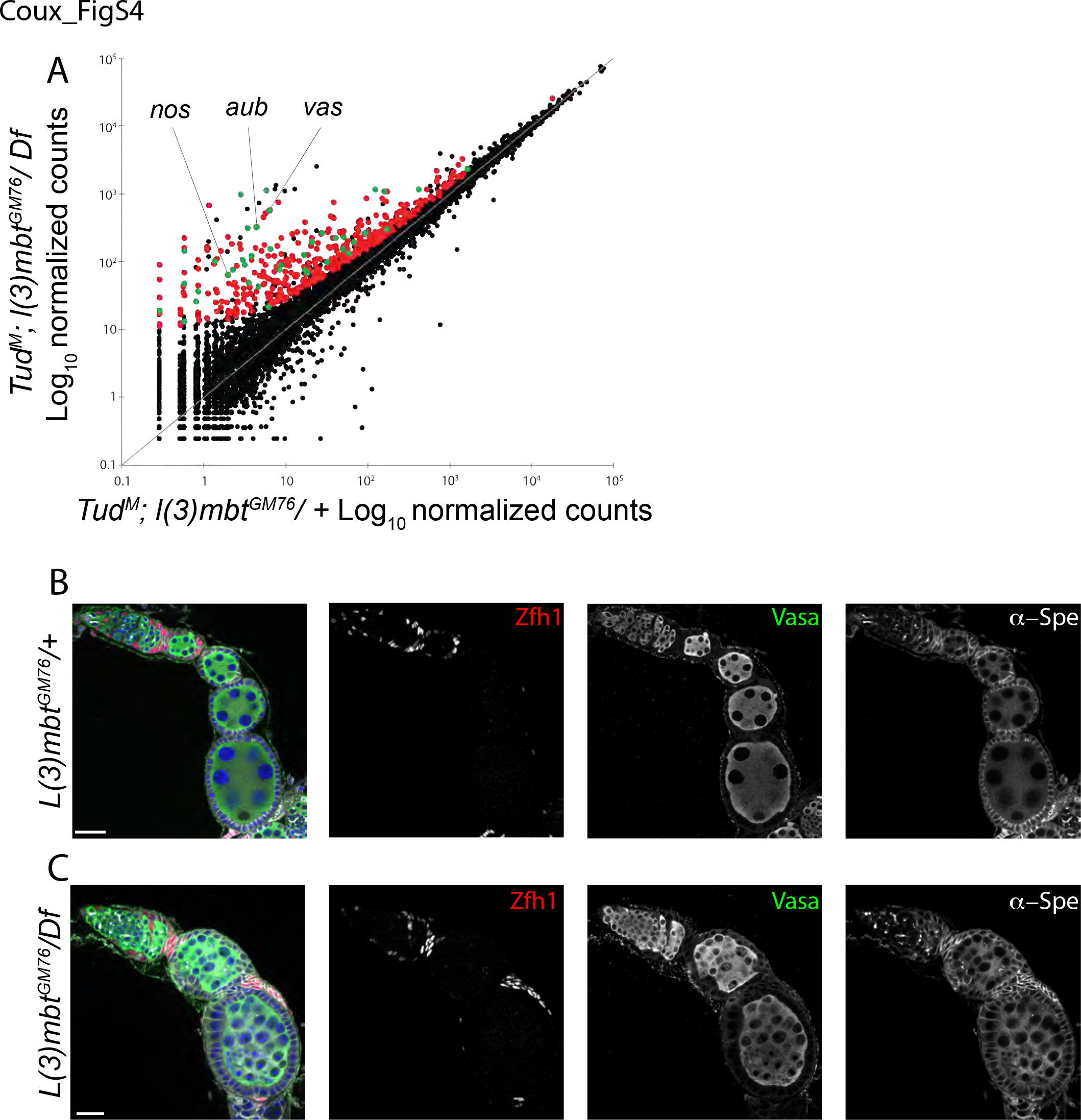
*L(3)mbt* mutant somatic cells are properly specified but ectopically express germline genes. (*A*) Scatterplot showing the expression of genes in *tud*^M^; *l(3)mbt*^*GM76*^/+ and *tud*^M^; *l(3)mbt*^*GM76*^ mutant ovaries, as measured by RNA-seq (normalized counts, log_10_). Derepressed genes are shown in red and de-repressed MBTS genes in green. (*B-C*) Confocal images of (*B*) control and *(C) l(3)mbt* mutant ovarioles stained for Vasa (green), Zfh1 (red), α-Spectrin (grey), and DAPI (blue). (*B*) In control ovaries, Zfh1 is expressed in escort cells, pre-follicle cells as well as stalk cells, which separate egg chambers. *(C) L(3)mbt* mutant ovariole showing normal Zfh1 expression but stalk cells accumulate on top of follicle cells. Scale bars, 25 μm.

**Supplemental figure 5.**
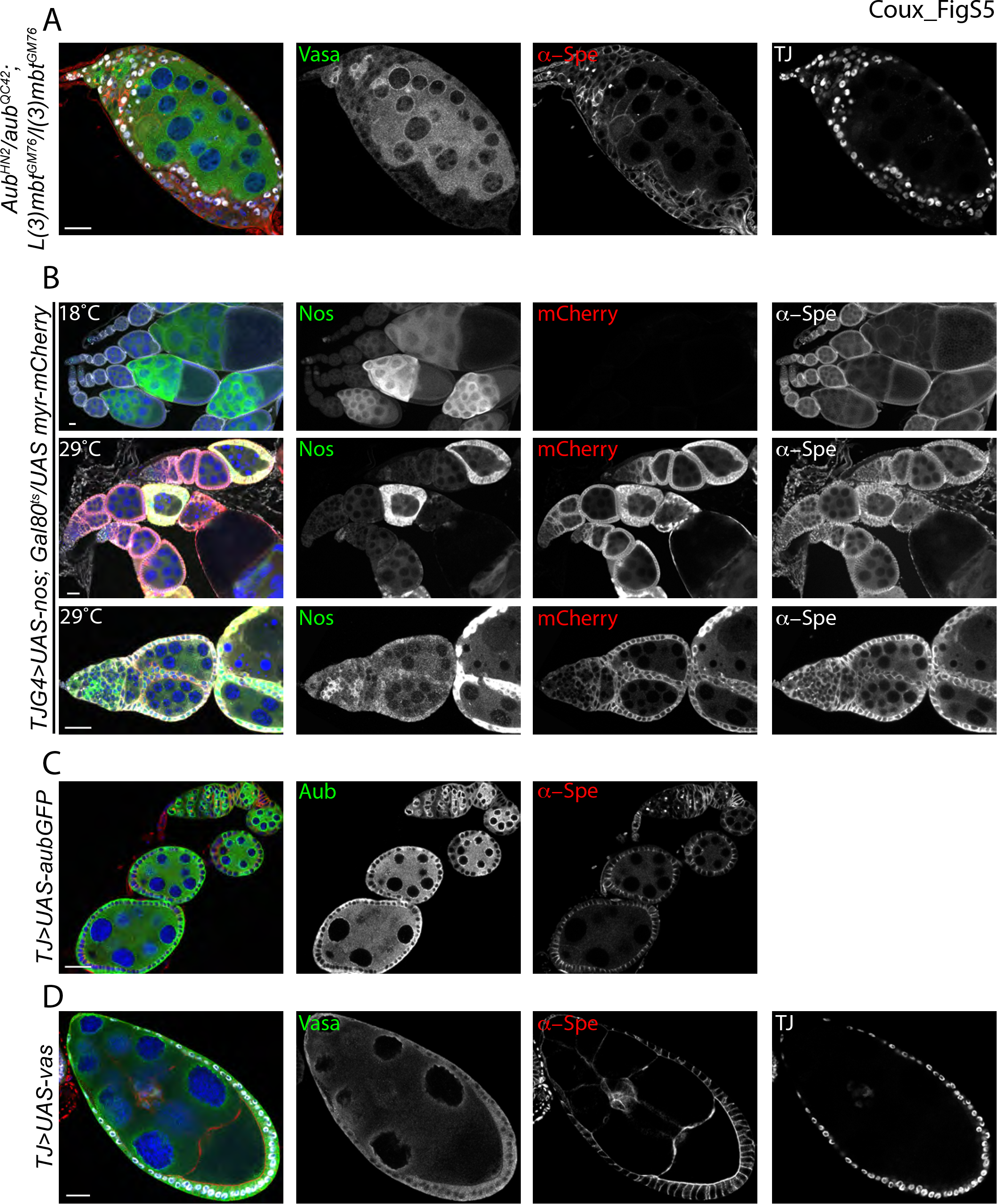
*Nos* but not *aub* or *vas* ectopic expression interferes with normal ovarian development. *(A)* Confocal image of *aub*; *l(3)mbt* double mutant ovariole stained for Vasa (green), α-Spectrin (red); TJ (grey), and DAPI (blue). The phenotype is similar to single *l(3)mbt* mutant. *(B)* Confocal images of ovaries expressing *UAS-nos*; *UAS-myr-mCherry* transgenes in somatic cells using a temperature sensitive system (Gal80^ts^) to express *nos* only in the adult. Ovaries were stained for Nos (green), mCherry (red), α-Spectrin (grey), and DAPI (blue). At 18°C, the transgenes are not expressed and ovarioles develop normally. When shifted at 29°C after eclosion, ectopic Nos expression in somatic cells perturbs egg chamber individualization and causes cell death. *(C-D)* Confocal images of ovaries misexpressing Aub or Vasa in somatic cells using the TJ-Gal4 driver and the (*C*) UAS-aubGFP and (*D*) UAS-vas transgenes. Ovaries were stained for (*A*) Aub (green), α-Spectrin (red), and DAPI (blue), or (*D*) Vasa (green), α-Spectrin (red), and TJ (grey). Scale bars, 25 μm.

**Supplemental figure 6.**
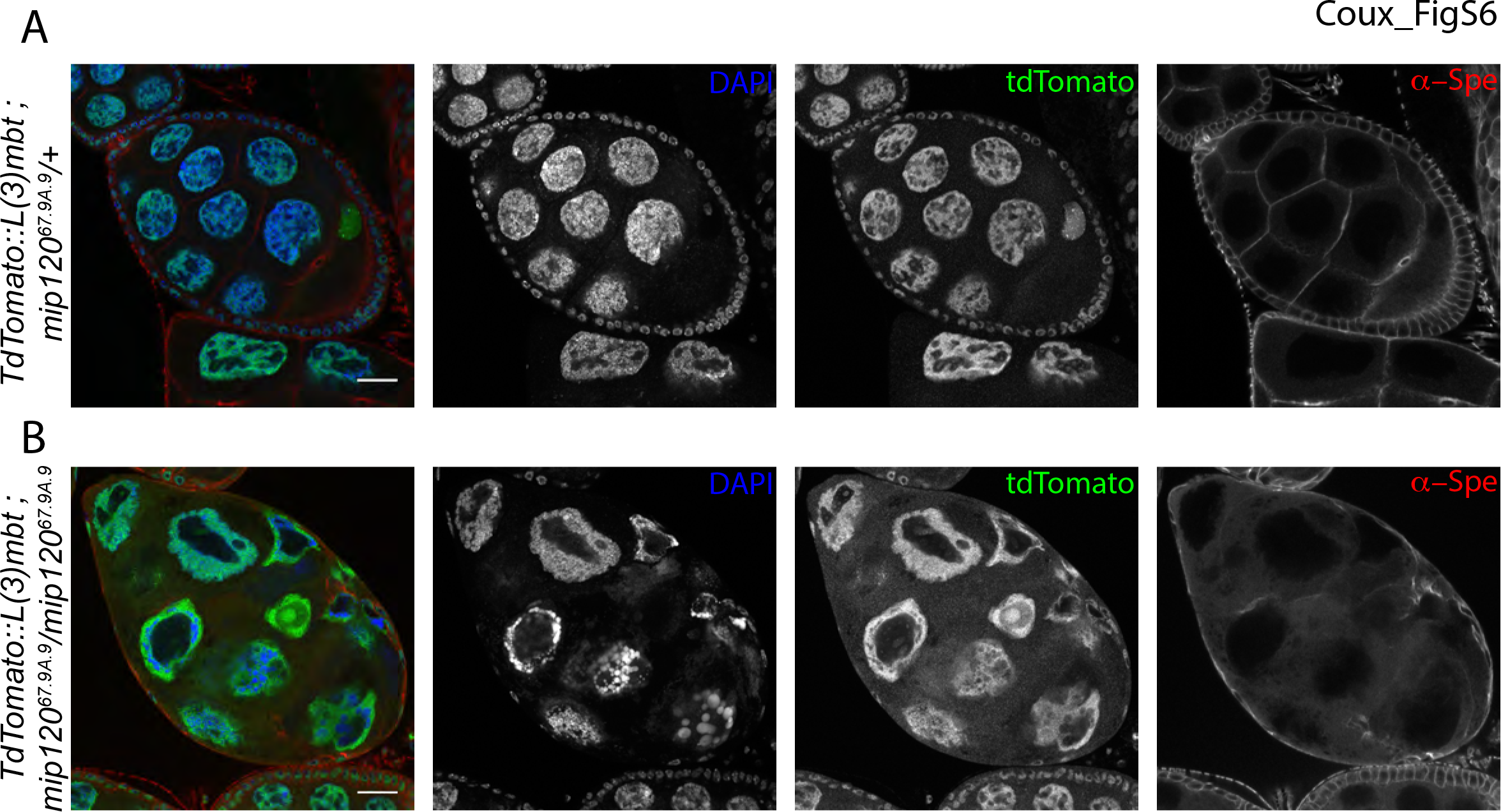
L(3)mbt nuclear localization is not affected in in *mip120*^*67.9A.9*^ mutant ovaries. *(A-B)* Confocal images of control and *mip120* mutant ovaries expressing the tdTomato::L(3)mbt fusion and stained for α-Spectrin (red), tdTomato (green), and DAPI (blue). TdTomato::L(3)mbt is nuclear and colocalizes with DAPI in both control and mutant ovaries. Mutant nurse cells nuclei are highly vacuolated. Scale bars, 25 μm.

**Supplemental figure 7.**
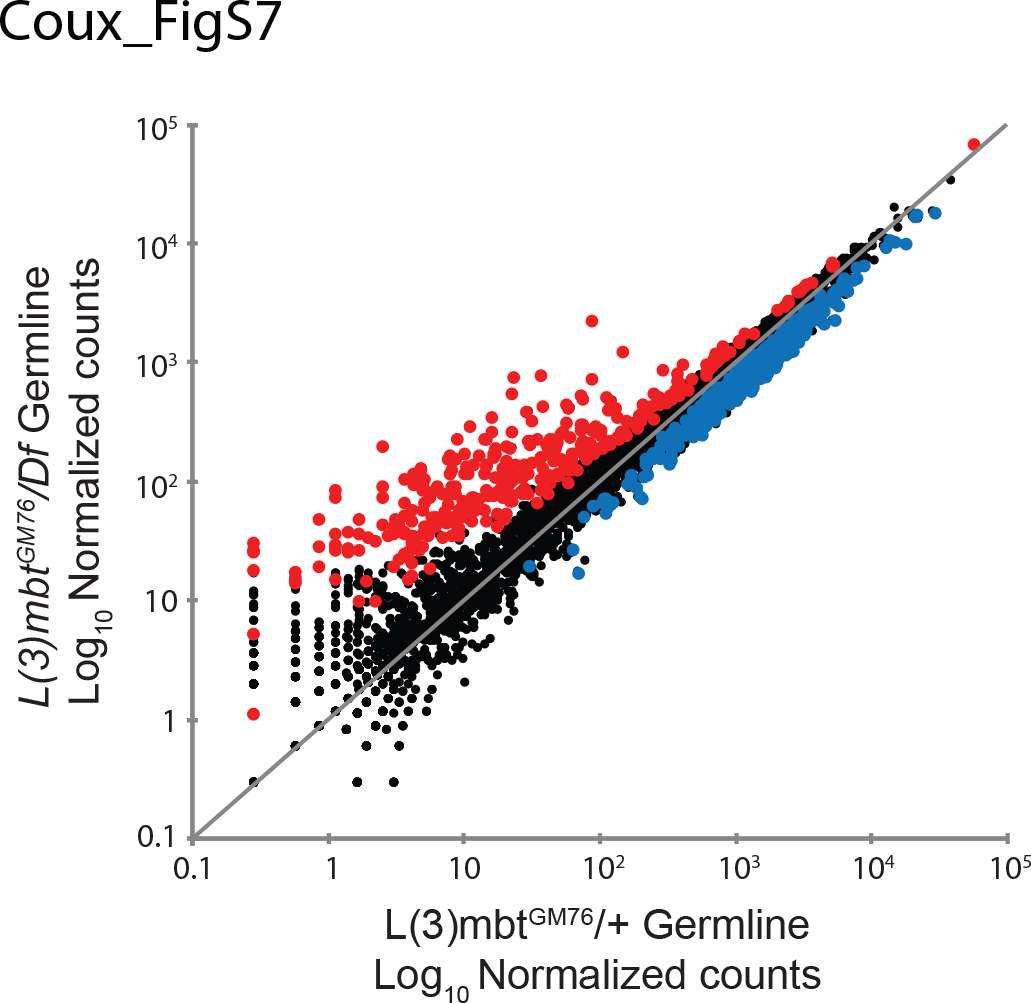
Scatterplots showing expression of genes in control and *l(3)mbt*^*GM76*^ mutant early embryos, as measured by RNA-seq (normalized counts, log_10_). Up-regulated genes are shown in red, down-regulated genes are shown in blue.

## Material and methods

### Fly stocks

*FRT82B*, *l(3)mbt^GM76^*, *e*/*TM6b* was generated in the Lehmann lab (Yohn et al., 2003) and secondary mutations removed (Richter et al., 2011). The following stocks were obtained from the Kyoto Stock Center:*w^1118^*;; *Df(3R)ED19066*/*TM6c* (#150208) and y* *w*\*;*P{GawB}NP1624*/*CyO*, *P*{UAS-lacZ.UW14}UW14 (*Tj*-*Gal4*, #104055); from Bloomington Drosophila Stock Center: *w1118; P{neoFRT}82B P{Ubi-GFP(S65T)nls}3R/TM6B*, *Tb*^*1*^ (BDSC #32655), *y*^*1*^ *w*\*; *P{UAS-FLP.D}JD1 (BDSC #4539)*, and *w*\*; *P{tubP-GAL80*^*ts*^*}2/TM2* (BDSC #7017). The following transgenes *C587-Gal4*, *UAS-nos-tub* (Clark et al., 2002; Ye et al., 2004), *UAS-vas* (Sengoku et al., 2006), *UASp-mCherry-myr*, *UAS-Aub-GFP* (Harris and Macdonald, 2001), and *TdTomato::l(3)mbt* (Blanchard et al., 2014) were obtained from the Xie, Jan, Nakamura, Zallen, Macdonald, and Botchan labs, respectively. The *mip120*^*67.9A.9*^ (Beall et al., 2007) and *FRT40A e2f2*^c03344^ (Ambrus et al., 2007) mutations were generated by the Botchan and Frolov labs. The following mutations are from the Lehmann lab stocks: *tud*^*1*^, *tud*^*B42*^ (Arkov et al., 2006), *aub*^*HN2*^, *aub*^*QC42*^(Schupbach and Wieschaus, 1991), *nos*^*L7*^ (Wang and Lehmann, 1991), and nos^B^ (Wang et al., 1994). All stocks were maintained at 18°C and crosses were performed at 25°C unless otherwise stated.

### Generation of transgenic lines

To generate *UASp*-*l(3)mbt::myc* transgenic flies, *l(3)mbt* coding sequence was amplified from LD05287 gold cDNA (Drosophila Gemonics Resource Center), cloned using the p-ENTR/D-TOPO system and recombined into the pPWM destination vector (Drosophila Gateway Vector Collection) using Gateway technology (Invitrogen). *pPWM*-*l(3)mbt* was randomly inserted on the 2^nd^ chromosome through P-transposition. To generate the *lint1* mutation, the target sequence (chrX:11044844-11044866) was identified by the flyCRISPR Optimal Target Finder tool (Gratz et al., 2014), amplified from genomic DNA using *lint*-*CRISPR* oligos (Supplemental Oligos Table) and ligated in the pU6-BbsI-gRNA plasmid (Gratz et al., 2013). The resulting construct was injected in *FRT19A;; vas*-*Cas9* embryos and progeny was screened by PCR and sequencing.

### Immunofluorescence

Adult ovaries from 2-3 day-old fattened females were dissected in cold PBS and fixed in 4% PFA for 20 min. Ovaries were permeabilized with 0.2% Triton X-100 in PBS and blocked with 1% (w/v) bovine serum albumin (BSA) and 0.2% Triton (PBST). Samples were incubated with primary antibodies in PBST overnight at 4°C. Next, ovaries were washed and incubated with secondary antibodies in PBST for 2 hours at room temperature. After three washes in PBS-0.2% Triton for 20 min each (including one containing 1:1000 DAPI), ovaries were mounted in SlowFade^®^ Gold mountant (Invitrogen) and imaged on Zeiss LSM780 or 800 confocal microscopes using 10, 20 or 43x objectives. Larval ovaries were processed the same way but permeabilized for 4 to 8 hours prior to primary antibody incubation. The following primary antibodies were used: rabbit α-Vasa (1:5000, Lehmann lab); goat α-Vasa (1:200, Santa Cruz Biotechnology sc-26877); mouse α-spectrin (1:200, DSHB); mouse α-Orb (4H8, 1:200, DHSB); chicken α-GFP (1:1000, Aves GFP-1020); goat α-tj (1:7000, kind gift of Dorothea Godt (Li et al., 2003)); rabbit α-zfh1 (1:5000, Lehmann lab); rat α-RFP (1:500, Chromotek 5F8); rabbit α-Nanos (1:200, kind gift of Prof. Nakamura); rabbit α-Aub (1:1000, Lehmann lab); rabbit α-DsRed (1:500, Living Colors *#*632496). Alexa Fluor 647 Phalloidin (1:500 Life Technologies) and rabbit α-myc Alexa fluor 555 conjugated (Millipore 16-225) were used as secondary antibodies. Alexa Fluor 488- (Life Technologies), Cy3- or Alexa Fluor 647-conjugated (Jackson Immunoresearch) secondary antibodies were used at a 1:1000 dilution.

### RNA sequencing

60-70 ovaries from females of maternal *tud^1^*/*tud*^*B42*^ and zygotic *l(3)mbt*^*GM76*^/+ or *l(3)mbt*^*GM76*^/*Df* genotypes (see Supplemental File 1) were dissected in cold PBS and RNA was extracted using TRIzol (Invitrogen) following the manufacturer’s protocol. For early embryos, *TJ*>*UAS*-*l(3)mbt::myc;l(3)mbt*^*GM76*^/+ and *TJ*>*UAS*-*l(3)mbt::myc;l(3)mbt*^*GM76*^/*Df* females were allowed to lay for 30 minutes to 1 hour on agar plates. Embryos were dechorionated in 50% bleach for 5 minutes, rinsed with PBS, and then lysed in TRIzol. Libraries were generated from 1μg of total RNA using the NEBNext Poly(A) magnetic Isolation Module (NEB *#*7490) and the NEBNext Ultra Directional RNA Library Prep Kit for Illumina (NEB *#* E7420). Libraries from biological replicates (three for ovaries, two for embryos) were sequenced on an Illumina Hi-Seq2000, single-end 50 run.

### RNA-seq data analysis

Sequencing results were demultiplexed and converted to FASTQ format using Illumina Bcl2FastQ software (Illumina). Reads were aligned to the fly genome (build dm3/BDGP5) using the splice-aware STAR aligner (Dobin et al., 2013). PCR duplicates were removed using the Picard toolkit (https://broadinstitute.github.io/picard/). HTSeq package was used to generate counts for each gene based on how many aligned reads overlap its exons (Anders et al., 2015). These counts were then used to test for differential expression using negative binomial generalized linear models implemented by the DESeq2 R package.

### Tissue expression clustering

Expression of deregulated genes was extracted from FlyAtlas (Chintapalli et al., 2007) using FlyBaseIDs, normalized to fly average and log2 transformed. Distance matrix was calculated using the “Manhattan” method and data clustered using “ward.D2”. Heatmap was generated using the heatmap.2 function of the gplots R package.

## Acknowledgements

We thank the entire Lehmann lab for discussion and input; C. Desplan, D. Keefe, E. Mazzoni, N. Dyson and J. Treisman for discussions and advice. We would like to thank J. Knoblich and C. Richter, M. Botchan and D. Blanchard, M.V. Frolov, D. Godt, S. Kobayashi and A. Nakamura for sharing reagents. We are grateful to the Bloomington Drosophila Stock Center (NIH P40OD018537) for fly strains; the Developmental Studies Hybridoma Bank, created by the NICHD of the NIH and maintained at The University of Iowa for antibodies and the NYUMC Genome Technology Center (NIH P30CA016087), for sequencing. R.X.C was supported by the New York State Stem Cell Program (NYSTEM) of the New York State Health Department (*#*C026880) and FKT was funded by EMBO and HFSP fellowships. RL is supported by NIH R37HD41900 and is an HHMI investigator.

## Author Contributions

R.X.C. and R.L. conceived the project. R.X.C. F.K.T. and R.L. designed experiments and wrote the manuscript. R.X.C. performed the experiments analyzed the data.

## References

Ambrus, A. M., Nicolay, B. N., Rasheva, V. I., Suckling, R. J. and Frolov, M. V. (2007). dE2F2-independent rescue of proliferation in cells lacking an activator dE2F1. Mol Cell Biol 27, 8561–8570.

Anders, S., Pyl, P. T. and Huber, W. (2015). HTSeq--a Python framework to work with high-throughput sequencing data. Bioinformatics 31, 166–169.

Arkov, A. L., Wang, J.-Y. S., Ramos, A. and Lehmann, R. (2006). The role of Tudor domains in germline development and polar granule architecture. Development 133, 4053–4062.

Beall, E. L., Lewis, P. W., Bell, M., Rocha, M., Jones, D. L. and Botchan, M. R. (2007). Discovery of tMAC: a Drosophila testis-specific meiotic arrest complex paralogous to Myb-Muv B. Genes & Development 21, 904–919.

Blanchard, D. P., Georlette, D., Antoszewski, L. and Botchan, M. R. (2014). Chromatin reader L(3)mbt requires the Myb-MuvB/DREAM transcriptional regulatory complex for chromosomal recruitment. Proc Natl Acad Sci USA.

Bonasio, R., Lecona, E. and Reinberg, D. (2010). MBT domain proteins in development and disease. Semin Cell Dev Biol 21, 221–230.

Cayirlioglu, P., Bonnette, P. C., Dickson, M. R. and Duronio, R. J. (2001). Drosophila E2f2 promotes the conversion from genomic DNA replication to gene amplification in ovarian follicle cells. Development 128, 5085–5098.

Cheng, M.-H., Andrejka, L., Vorster, P. J., Hinman, A. and Lipsick, J. S. (2017). The Drosophila LIN54 homolog Mip120 controls two aspects of oogenesis. Biol Open 6, 967–978.

Chintapalli, V. R., Wang, J. and Dow, J. A. T. (2007). Using FlyAtlas to identify better Drosophila melanogaster models of human disease. Nat Genet 39, 715–720.

Christerson, L. B. and McKearin, D. M. (1994). orb is required for anteroposterior and dorsoventral patterning during Drosophila oogenesis. Genes & Development 8, 614–628.

Clark, I. E., Dobi, K. C., Duchow, H. K., Vlasak, A. N. and Gavis, E. R. (2002). A common translational control mechanism functions in axial patterning and neuroendocrine signaling in Drosophila. Development 129, 3325–3334.

Dobin, A., Davis, C. A., Schlesinger, F., Drenkow, J., Zaleski, C., Jha, S., Batut, P., Chaisson, M. and Gingeras, T. R. (2013). STAR: ultrafast universal RNA-seq aligner. Bioinformatics 29, 15–21.

Edgar, B. A. and Schubiger, G. (1986). Parameters controlling transcriptional activation during early Drosophila development. Cell 44, 871–877.

Gateff, E., Löffler, T. and Wismar, J. (1993). A temperature-sensitive brain tumor suppressor mutation of Drosophila melanogaster: developmental studies and molecular localization of the gene. Mech Dev 41, 15–31.

Georlette, D., Ahn, S., MacAlpine, D. M., Cheung, E., Lewis, P. W., Beall, E. L., Bell, S. P., Speed, T., Manak, J. R. and Botchan, M. R. (2007). Genomic profiling and expression studies reveal both positive and negative activities for the Drosophila Myb MuvB/dREAM complex in proliferating cells. 21, 2880–2896.

Gilboa, L. (2015). Organizing stem cell units in the Drosophila ovary. Curr Opin Genet Dev 32, 31–36.

Gilboa, L. and Lehmann, R. (2006). Soma-germline interactions coordinate homeostasis and growth in the Drosophila gonad. Nature 443, 97–100.

Godt, D. and Laski, F. A. (1995). Mechanisms of cell rearrangement and cell recruitment in Drosophila ovary morphogenesis and the requirement of bric à brac. Development 121, 173–187.

Gratz, S. J., Cummings, A. M., Nguyen, J. N., Hamm, D. C., Donohue, L. K., Harrison, M. M., Wildonger, J. and O’Connor-Giles, K. M. (2013). Genome engineering of Drosophila with the CRISPR RNA-guided Cas9 nuclease. Genetics 194, 1029–1035.

Gratz, S. J., Ukken, F. P., Rubinstein, C. D., Thiede, G., Donohue, L. K., Cummings, A. M. and O’Connor-Giles, K. M. (2014). Highly specific and efficient CRISPR/Cas9-catalyzed homology-directed repair in Drosophila. Genetics 196, 961–971.

Handler, D., Meixner, K., Pizka, M., Lauss, K., Schmied, C., Gruber, F. S. and Brennecke, J. (2013). The genetic makeup of the Drosophila piRNA pathway. Mol Cell 50, 762–777.

Harris, A. N. and Macdonald, P. M. (2001). Aubergine encodes a Drosophila polar granule component required for pole cell formation and related to eIF2C. Development 128, 2823–2832.

Harrison, D. A. and Perrimon, N. (1993). Simple and efficient generation of marked clones in Drosophila. Current Biology 3, 424–433.

Hayashi, Y., Hayashi, M. and Kobayashi, S. (2004). Nanos suppresses somatic cell fate in Drosophila germ line. Proc Natl Acad Sci USA 101, 10338–10342.

Hinz, U., Giebel, B. and Campos-Ortega, J. A. (1994). The basic-helix-loop-helix domain of Drosophila lethal of scute protein is sufficient for proneural function and activates neurogenic genes. Cell 76, 77–87.

Huynh, J. R. (2006). Fusome as a cell-cell communication channel. Cell-Cell Channels. Springer.

Huynh, J.-R. and St Johnston, D. (2004). The origin of asymmetry: early polarisation of the Drosophila germline cyst and oocyte. Current Biology 14, R438–49.

Janic, A., Mendizabal, L., Llamazares, S., Rossell, D. and Gonzalez, C. (2010). Ectopic Expression of Germline Genes Drives Malignant Brain Tumor Growth in Drosophila. Science 330, 1824–1827.

Jarriault, S., Schwab, Y. and Greenwald, I. (2008). A Caenorhabditis elegans model for epithelial-neuronal transdifferentiation. Proc Natl Acad Sci USA 105, 3790–3795.

Leatherman, J. L. and Dinardo, S. (2008). Zfh-1 controls somatic stem cell self-renewal in the Drosophila testis and nonautonomously influences germline stem cell self-renewal. Cell Stem Cell 3, 44–54.

Lee C.Y.S., Lu T. and Seydoux G. (2017). Specification of the germline by Nanos-dependent down-regulation of the somatic synMuvB transcription factor LIN-15B. Preprint (BiorXiv).

Lewis, P. W., Beall, E. L., Fleischer, T. C., Georlette, D., Link, A. J. and Botchan, M. R. (2004). Identification of a Drosophila Myb-E2F2/RBF transcriptional repressor complex. Genes & Development 18, 2929–2940.

Li, M. A., Alls, J. D., Avancini, R. M., Koo, K. and Godt, D. (2003). The large Maf factor Traffic Jam controls gonad morphogenesis in Drosophila. Nat Cell Biol 5, 994–1000.

Maimon, I., Popliker, M. and Gilboa, L. (2014). Without children is required for Stat-mediated zfh1 transcription and for germline stem cell differentiation. Development 141, 2602–2610.

McGuire, S. E., Mao, Z. and Davis, R. L. (2004). Spatiotemporal gene expression targeting with the TARGET and gene-switch systems in Drosophila. Sci. STKE 2004, pl6–pl6.

Meier, K., Mathieu, E.-L., Finkernagel, F., Reuter, L. M., Scharfe, M., Doehlemann, G., Jarek, M. and Brehm, A. (2012). LINT, a novel dL(3)mbt-containing complex, represses malignant brain tumour signature genes. PLoS Genet 8, e1002676.

Miles, W. O., Korenjak, M., Griffiths, L. M., Dyer, M. A., Provero, P. and Dyson, N. J. (2014). Post-transcriptional gene expression control by NANOS is up-regulated and functionally important in pRb-deficient cells. 33, 2201–2215.

Petrella, L. N., Wang, W., Spike, C. A., Rechtsteiner, A., Reinke, V. and Strome, S. (2011). synMuv B proteins antagonize germline fate in the intestine and ensure C. elegans survival. Development 138, 1069–1079.

Richter, C., Oktaba, K., Steinmann, J. and Knoblich, J. A. (2011). The tumour suppressor L(3)mbt inhibits neuroepithelial proliferation and acts on insulator elements. Nat Cell Biol 13, 1029–1039.

Schupbach, T. and Wieschaus, E. (1991). Female sterile mutations on the second chromosome of Drosophila melanogaster. II. Mutations blocking oogenesis or altering egg morphology. Genetics 129, 1119–1136.

Sengoku, T., Nureki, O., Nakamura, A., Kobayashi, S. and Yokoyama, S. (2006). Structural basis for RNA unwinding by the DEAD-box protein Drosophila Vasa. Cell 125, 287–300.

Smendziuk, C. M., Messenberg, A., Vogl, A. W. and Tanentzapf, G. (2015). Bi-directional gap junction-mediated soma-germline communication is essential for spermatogenesis. Development 142, 2598–2609.

Sumiyoshi, T., Sato, K., Yamamoto, H., Iwasaki, Y. W., Siomi, H. and Siomi, M. C. (2016). Loss of l(3)mbt leads to acquisition of the ping-pong cycle in Drosophila ovarian somatic cells. Genes & Development 30, 1617–1622.

Tursun, B., Patel, T., Kratsios, P. and Hobert, O. (2011). Direct Conversion of C. elegans Germ Cells into Specific Neuron Types. Science 331, 304–308.

Van Bortle, K., Nichols, M. H., Li, L., Ong, C.-T., Takenaka, N., Qin, Z. S. and Corces, V. G. (2014). Insulator function and topological domain border strength scale with architectural protein occupancy. Genome Biol 15, R82.

Wang, C. and Lehmann, R. (1991). Nanos is the localized posterior determinant in Drosophila. Cell 66, 637–647.

Wang, C., Dickinson, L. K. and Lehmann, R. (1994). Genetics of nanos localization in Drosophila. Dev Dyn 199, 103–115.

Weidmann, C. A., Qiu, C., Arvola, R. M., Lou, T.-F., Killingsworth, J., Tanaka Campbell, Z. T., Hall, T. M. and Goldstrohm, A. C. (2016). Drosophila Nanos acts as a molecular clamp that modulates the RNA-binding and repression activities of Pumilio. Elife 5, e17096.

Ye, B., Petritsch, C., Clark, I. E., Gavis, E. R., Jan, L. Y. and Jan, Y. N. (2004). Nanos and Pumilio are essential for dendrite morphogenesis in Drosophila peripheral neurons. Curr Biol 14, 314–321.

Yohn, C. B., Pusateri, L., Barbosa, V. and Lehmann, R. (2003). l(3)malignant brain tumor and three novel genes are required for Drosophila germ-cell formation. Genetics 165, 1889–1900.

Zhu, C.-H. and Xie, T. (2003). Clonal expansion of ovarian germline stem cells during niche formation in Drosophila. Development 130, 2579–2588.

